# Senescence-inhibitory Δ133p53α counteracts accelerated ageing and mortality

**DOI:** 10.64898/2025.12.31.697195

**Authors:** Leo Yamada, Huaitian Liu, Natalia von Muhlinen, Curtis C. Harris, Izumi Horikawa

## Abstract

Research on progeria not only contributes to treatments for the disease but also enhances our understanding of physiological ageing^1^. Mouse models of progeria recapitulate pathological ageing phenotypes seen in patients, including cardiovascular defects, increased cellular senescence, systemic inflammation, DNA damage accumulation, and shortened lifespan^2^. In cultured cells from Hutchinson-Gilford progeria syndrome (HGPS) patients, the human p53 isoform Δ133p53α was previously shown to inhibit p53-mediated cellular senescence, proinflammatory IL-6 production, and DNA damage accumulation, and to extend cellular replicative lifespan^3^. Here we show that, in a heterozygous HGPS mouse model^4^, transgenic expression of Δ133p53α reproduces these *in vitro*-observed effects across multiple organs *in vivo* and extends median lifespan by 11% (387 versus 349 days, *P* = 0.0379). In the aorta and skin, Δ133p53α abrogates progeria-characteristic pathological changes and preserves tissue integrity. Our data further suggest that Δ133p53α may promote a broad spectrum of ageing-counteracting mechanisms, including bone homeostasis, metabolic fitness, antioxidant defense, youthful epigenome, and tissue stemness. Together with the anti-inflammatory and tissue-preserving effects of Δ133p53α in naturally aged mice and its age-associated downregulation in human tissues, this study suggests that Δ133p53α-based therapeutic strategies may be applicable not only to HGPS but also as broader interventions for preventing or delaying ageing.

---

Hutchinson-Gilford progeria syndrome (HGPS), a notable example of progeroid syndromes characterized by accelerated ageing symptoms^5^, is caused by mutations in the *LMNA* gene, which result in the production of a defective nuclear envelope protein known as progerin^6^. Although potentially promising therapies targeting the synthesis, modification, or clearance of progerin have been developed^5^, further efforts are critically needed to establish improved treatment regimens for patients with HGPS. Since some, though not all, ageing symptoms that emerge in childhood among HGPS patients also appear later in life during natural ageing^1^, cellular, molecular and phenotypic studies on this disease hold significant potential for advancing our understanding of natural ageing processes and identifying interventions. Among several mouse models of HGPS, *Lmna^G609G^* mice carry the mutant human *LMNA* allele (c.1827C>T;p.Gly609Gly) knocked-in at the mouse *Lmna* locus^4^, thereby reproducing the patient-derived, progerin-producing mutation in mice. These mice phenocopy the major clinical features of HGPS, including cardiovascular, skeletal and skin abnormalities, as well as shortened lifespan^4^.

The tumor suppressor protein p53 is a key contributor to both cellular ageing, referred to as cellular senescence, and organismal ageing, while it literally plays critical roles in tumor suppression^7,8^. When attempting to manipulate p53 or its upstream or downstream signaling for therapeutic purposes, it is therefore essential to consider and balance the beneficial and adverse aspects of p53 activities, for example, p53-mediated maintenance of genome stability and p53-mediated acceleration of cellular senescence and ageing, respectively^7,8^. Δ133p53α is a naturally occurring isoform of human p53, which lacks the N-terminal 132 amino acids due to alternative transcription initiation from intron 4 and alternative translation initiation at codon 133 methionine^9^. Missing the N-terminal transactivation domains but retaining the C-terminal oligomerization domain, Δ133p53α physically interacts with p53 and functions as a dominant-negative inhibitory isoform of p53^9,10^. Of note, the dominant-negative inhibition of Δ133p53α preferentially acts against p53-inducible genes involved in cellular senescence (for example, p21^WAF1/CIP1^, or CDKN1A) through a mechanism(s) yet to be fully elucidated, while the expressions of DNA repair genes are preserved^10,11^. In various types of normal human cells cultured *in vitro*, including fibroblasts^9^, CD8^+^ T lymphocytes^12^, prostate epithelial cells^13^, ectocervical keratinocytes^13^, and brain astrocytes^14,15^, retroviral or lentiviral expression of Δ133p53α reduces the expression of p21^WAF1/CIP1^ and delays the onset of replicative senescence, extending replicative lifespan, or mitigates exogenous stress-induced senescence. These findings suggest that Δ133p53α may have therapeutic values in diseases or dysfunctional conditions associated with high levels of cellular senescence, with minimal risks of genome instability due to deficient p53-mediated DNA repair.

In our previous study using *in vitro* cultured fibroblasts derived from HGPS patients^3^, Δ133p53α prevented these cells from early entry into replicative senescence, extended their replicative lifespan, and reduced the production of proinflammatory interleukin 6 (IL-6), as it did in normal cell types^9,12–15^. It also mitigated the progerin-induced accumulation of DNA damage^3^. In this study, by generating Δ133p53α-transgenic mice and crossbreeding them with the progeria model *Lmna^G609G^* mice^4^, we investigate the *in vivo* therapeutic effects of Δ133p53α on the accelerated ageing phenotypes characteristic of HGPS at the cellular, organ, and organismal levels. Our examination of Δ133p53α-transgenic mice on a wild-type background, along with analysis of Δ133p53α expression in human databases, suggests a role for Δ133p53α in natural ageing in mice and humans as well.

## Δ133p53α-transgenic mice

Using the ROSA26 knock-in vector engineered for Cre/loxP-inducible transgenic expression of Δ133p53α (Extended Data Fig. 1a), we successfully generated a mouse strain carrying the expected allele (CAG-LSL-Δ133p53α) at the *ROSA26* locus (Extended Data Fig. 1b). The mice heterozygous for this knocked-in allele (termed *CAG-133^LSL/+^*) were crossed with heterozygous UBC-Cre-ERT2 mice ubiquitously expressing a tamoxifen-activated Cre recombinase (termed *Cre^Tg/+^*) to obtain *CAG-133^LSL/+^*;*Cre^Tg/+^* mouse embryonic fibroblasts (MEFs). These MEFs, along with control *CAG-133^+/+^*;*Cre^+/+^* MEFs, were used in a series of experiments to verify that Δ133p53α, which is a human sequence, functions in mouse cells (Extended Data Fig. 2). Treatment with 4-hydroxytamoxifen (4-OHT), best at 1.0 µM for 72 h, resulted in an induction of Δ133p53α in *CAG-133^LSL/+^*;*Cre^Tg/+^* MEFs (Extended Data Fig. 2a). The induced expression of Δ133p53α inhibited proliferation arrest and extended replicative lifespan (Extended Data Fig. 2b), reduced the number of SA-β-gal-positive senescent cells (Extended Data Fig. 2c), suppressed the senescence-associated increase in p21^Waf1/Cip1^ (Cdkn1a) expression (Extended Data Fig. 2d), and alleviated DNA damage, as indicated by γ-H2AX foci (Extended Data Fig. 2e). These phenotypes recapitulate those observed with vector-driven expression of Δ133p53α in various human cell types^3,9,12–15^. The functional compatibility of Δ133p53α in mouse cells provides the basis for this present study to investigate its *in vivo* effects in mice.

To confirm tamoxifen-inducible expression of Δ133p53α in mice *in vivo*, two *CAG-133^LSL/+^*;*Cre^Tg/+^* mice (from *CAG-133^LSL/+^* × *Cre^Tg/+^* crossing above) were intraperitoneally (i.p.) injected with tamoxifen on five consecutive days at 8-9 weeks of age and then euthanized at 15 weeks of age, along with a *CAG-133^LSL/+^*;*Cre^Tg/+^*mouse that received no tamoxifen and a wild-type *CAG-133^+/+^*;*Cre^+/+^* mouse. In Western blot analysis, all tissues examined (brain, skin, liver, lung, aorta, thymus and spleen) from the two tamoxifen-injected *CAG-133^LSL/+^*;*Cre^Tg/+^*mice, but not from non-injected and wild-type controls, showed induced expression of Δ133p53α (Extended Data Fig. 3), confirming the widespread induction of Δ133p53α in this model with the ubiquitous expression of tamoxifen-activated Cre recombinase. Hereafter in this study, we designate all tamoxifen-injected *CAG-133^LSL/+^*;*Cre^Tg/+^* mice, across wild-type, heterozygous and homozygous *Lmna^G609G^* backgrounds, as *CAG-133^Tam/+^*;*Cre^Tg/+^* mice (Group 1) to distinguish them from non-injected *CAG-133^LSL/+^*;*Cre^Tg/+^*mice on wild-type and heterozygous *Lmna^G609G^* backgrounds (Group 2). This study also includes two other control groups: tamoxifen-injected *CAG-133^+/+^*;*Cre^Tg/+^* mice on heterozygous and homozygous *Lmna^G609G^* backgrounds (Group 3); and non-injected *CAG-133^+/+^*;*Cre^+/+^* mice on all three backgrounds (Group 4).

### Δ133p53α effects recapitulated *in vivo*

To examine whether transgenic expression of Δ133p53α affects ageing-associated cellular phenotypes *in vivo*, *CAG-133^LSL/+^*;*Cre^Tg/+^*mice were crossed with heterozygous *Lmna^G609G^* mice (termed *Lmna^G609G/+^*) to obtain eight *CAG-133^LSL/+^*;*Cre^Tg/+^*;*Lmna^G609G/+^*mice. Four of them were randomly selected, injected i.p. with tamoxifen on five consecutive days at 5-6 weeks of age, and euthanized at 15 weeks of age (Group 1, *CAG-133^Tam/+^*;*Cre^Tg/+^*;*Lmna^G609G/+^*), along with four non-injected counterparts (Group 2, *CAG-133^LSL/+^*;*Cre^Tg/+^*;*Lmna^G609G/+^*) and four age-matched *CAG-133^+/+^*;*Cre^+/+^*;*Lmna^G609G/+^*mice (Group 4). Two anti-p53 antibodies, sheep polyclonal SAPU and mouse monoclonal DO-11, which theoretically recognize both full-length p53 and Δ133p53α, were first tested on the same Western blot membrane, directing us to primarily use SAPU for detecting Δ133p53α and DO-11 for full-length p53 in this study (Extended Fig. 4a,b). In seven tissues examined, including the skin (dorsal back), skeletal muscle (femoris muscle), kidney, spleen, lung, heart, and brain, all four Group-1 mice, but not Group-2 or-4 mice, showed transgenic expression of Δ133p53α, accompanied by decreased, increased or unchanged expression of full-length p53 (Fig. 1a-e and Extended Fig. 4c,d). In the skin, muscle, kidney, spleen, and lung (Fig. 1a-e), the Group-1 mice exhibited significantly decreased expression of p21^Waf1/Cip1^, a p53-inducible gene involved in cellular senescence^16^ and a recommended *in vivo* marker of senescence^17^, although its expression levels were too low to quantify in the heart and brain (Extended Fig. 4c,d). Decreased SA-β-gal positivity was also observed in the spleen of Group-1 mice compared to Group-2 and-4 mice (Extended Data Fig. 4e). Among the six tissues in which progerin levels could be quantified, three of them (skin, muscle, and kidney) showed no significant change upon Δ133p53α expression (Fig. 1a-c), whereas progerin levels were increased with Δ133p53α expression in the heart (Extended Data Fig. 4c) and decreased in the spleen and lung, significantly and non-significantly, respectively (Fig. 1d,e).

**Fig. 1.**
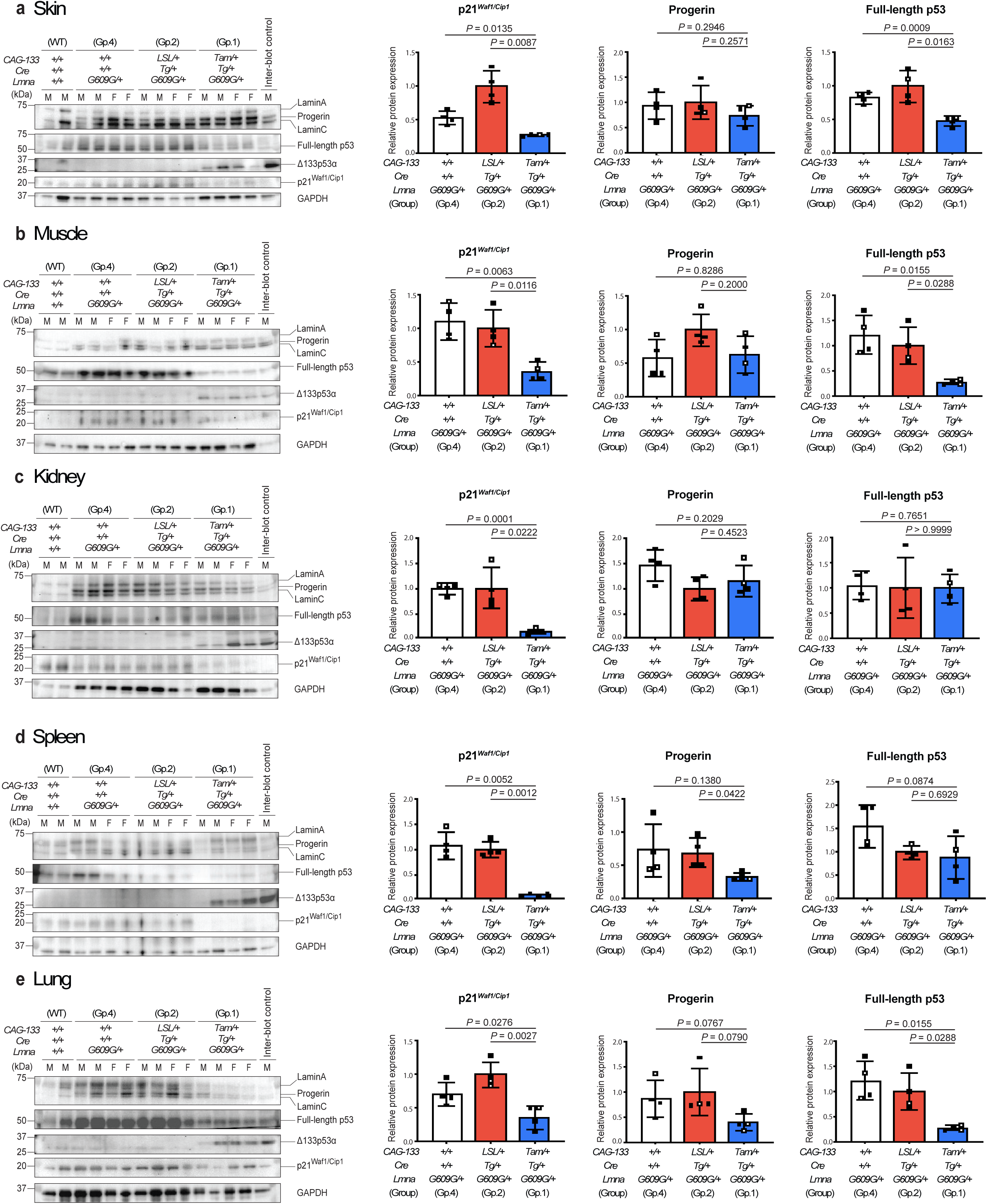
Transgenic expression of Δ133p53α represses p21^Waf1/Cip1^ (Cdkn1a) in heterozygous *Lmna^G609G/+^* progeria model mice. Western blot analyses of lamin A/C and progerin, full-length p53, Δ133p53α, and p21^Waf1/Cip1^ were performed in four 15-week-old Δ133p53α-expressing Group-1 mice (*CAG-133^Tam/+^*;*Cre^Tg/+^*;*Lmna^G609G/+^*), along with four each of age-matched, non-expressing control Group-2 (*CAG-133^LSL/+^*;*Cre^Tg/+^*;*Lmna^G609G/+^*) and Group-4 mice (*CAG-133^+/+^*;*Cre^+/+^*;*Lmna^G609G/+^*), as well as two age-matched wild-type mice (*CAG-133^+/+^*;*Cre^+/+^*;*Lmna^+/+^*). Results from skin (**a**), skeletal muscle (**b**), kidney (**c**), spleen (**d**), and lung (**e**) are presented. An inter-blot control (liver from a Group-1 mouse) was included in all blots. F, female; M, male. GAPDH was a loading control and used for normalization of p21^Waf1/Cip1^, progerin, and full-length p53 expression levels. Quantitative data summaries of p21^Waf1/Cip1^, progerin, and full-length p53 in Group-1,-2 and-4 mice are shown as relative values to Group-2 mice (mean ± s.d. from n = 4; open circles indicate two females, and closed circles indicate two males). *P* values were determined by Welch’s *t*-test. Two wild-type mice were used only as references and not for statistical comparisons.

To obtain further evidence for Δ133p53α-induced cellular changes, the aortic medial layer (tunica media) and the skin, which contain vascular smooth muscle cells (VSMCs) and hair follicle cells, respectively, two major cell types affected in the *Lmna^G609G^* model^2^, were examined in immunohistochemical (IHC) staining (Fig. 2a-f). The percentage of p21^Waf1/Cip1^-positive nuclei was significantly lower in both tissues in Group-1 mice than in those in Group-2 and Group-4 controls (Fig. 2a,d), consistent with the results shown in Fig. 1. p16^Ink4a^ (Cdkn2a), another *in vivo* marker of senescence^17^, also showed significantly lower positivity in Group-1 mice compared with Group-2 and Group-4 controls (Fig. 2b,e). Progerin-induced accumulation of DNA damage^1^, reflected by an ageing-associated increase in γ-H2AX-positive nuclei^18^, was significantly reduced in Group-1 mice compared with Group-2 and Group-4 controls (Fig. 2c,f). The reduced numbers of p21^Waf1/Cip1^-, p16^Ink4a^-, and γ-H2AX-positive nuclei in Group-1 mice were observed in the skeletal muscle as well (Extended Data Fig. 5a-c).

**Fig. 2.**
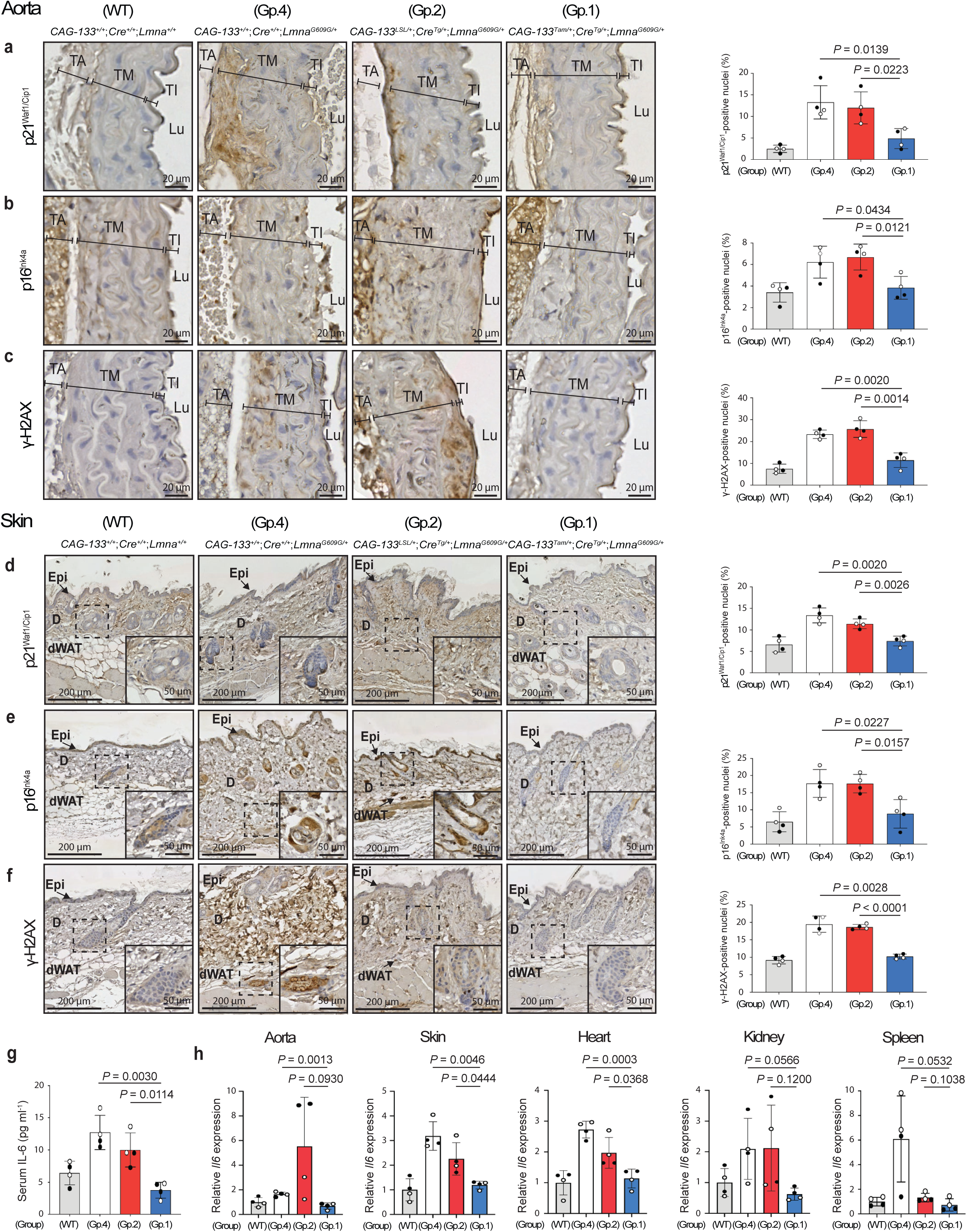
Δ133p53α reduces cellular senescence, DNA damage, and pro-inflammatory IL-6. The tissue section slides of the aorta (**a-c**) and skin (**d-f**) were prepared from the same set of Group-1, Group-2 and Group-4 mice as in Fig. 1, as well as four age-matched wild-type mice (the same two as in Fig. 1 and the additional two), followed by immunohistochemical (IHC) staining of p21^Waf1/Cip1^ (**a,d**), p16^Ink4a^ (**b,e**), and γ-H2AX (**c,f**). Representative images are shown on the left (scale bars, 20 µm in **a-c** and 200 µm in **d-f**). Insets in **d-f** show enlarged images of the boxed areas containing hair follicles (scale bars, 50 µm). TA, tunica adventitia; TM, tunica media; TI, tunica intima; Lu, Lumen; Epi, epidermis; D, dermis; dWAT, dermal white adipose tissue. Positively stained nuclei were counted in the tunica media containing vascular smooth muscle cells (VSMCs) (**a-c**) and in the dermis containing fibroblasts and hair follicles (**d-f**). Quantitative summaries are presented on the right (mean ± s.d. from n = 4; open circles, females; closed circles, males). *P* values were determined by Welch’s *t*-test. **g,** Serum IL-6 concentrations (pg ml^-1^) quantitated by ELISA in the same set of mice as above (mean ± s.d. from n = 4; open circles, females; closed circles, males). *P* values were calculated by Welch’s *t*-test. **h,** Il6 mRNA levels quantitated by qRT-PCR in the aorta, skin, heart, kidney, and spleen from the same set of mice as above (mean ± s.d. from n = 4; each with technical triplicate). *P* values were calculated by Welch’s *t*-test. Muscle and lung tissues were also tested, but the Ct values were not sufficiently high for reliable analysis.

Serum levels of the proinflammatory cytokine IL-6, a component of the senescence-associated secretory phenotype and a mediator of systemic and local inflammatory states^19^, were significantly lower in Group-1 mice than in Group-2 and Group-4 controls, being comparable to the low level observed in age-matched wild-type mice (Fig. 2g). The aorta, skin, heart, kidney, and spleen of Group-1 mice, compared with those of Group-2 and Group-4 controls, also showed lower levels of Il6 mRNA expression (with statistically significant reductions relative to one or both controls in the former three tissues, and borderline or non-significant trends in the latter two), reaching levels comparable to wild-type mice (Fig. 2h). In summary, transgenic expression of Δ133p53α *in vivo* systemically recapitulates its cellular effects observed *in vitro*, counteracting cellular senescence, DNA damage accumulation, and IL-6 production.

### Δ133p53α mitigates progeria phenotypes

To examine whether Δ133p53α can ameliorate progeria-associated pathological changes and shortened lifespan in heterozygous *Lmna^G609G^*mice, we generated an additional set of *Lmna^G609G/+^* groups as follows: Group 1, *CAG-133^Tam/+^*;*Cre^Tg/+^*;*Lmna^G609G/+^*(*CAG-133^LSL/+^*;*Cre^Tg/+^*;*Lmna^G609G/+^*that received tamoxifen at 5-6 weeks of age, n = 5 for scheduled euthanasia and n = 15 for survival analysis); Group 3, *CAG-133^+/+^*;*Cre^Tg/+^*;*Lmna^G609G/+^*, which received tamoxifen at 5-6 weeks of age (n = 5 for scheduled euthanasia and n = 17 for survival analysis); and Group 4, *CAG-133^+/+^*;*Cre^+/+^*;*Lmna^G609G/+^* without tamoxifen injection (n = 16 for survival analysis). The scheduled euthanasia of Group-1 and Group-3 mice (n = 5 each) was performed at 9-10 months of age, along with age-matched wild-type mice (n = 5), following CT imaging to examine spinal kyphosis. All other mice were monitored throughout their lifespan.

In tissue sections of the aorta (aortic arch) and skin (dorsal back) from these 9-10-month-old mice, the numbers of p21^Waf1/Cip1^-, p16^Ink4a^-, and γ-H2AX-positive cells were reduced in Group-1 mice compared with Group-3 mice, reaching levels comparable or close to those in wild-type mice (Extended Data Fig. 6a,b). These findings reproduce the earlier results that compared Group-1 and Group-2 mice at 15 weeks of age (Fig. 2). In the aorta, the tunica media is the main affected layer^2^ and was identified by the presence of elastic fibers (Fig. 3a). The progeria-characteristic reduction of cell numbers in the tunica media (largely VSMCs) was evident in Group-3 mice, but was restored to wild-type levels in Δ133p53α-expressing Group-1 mice (Fig. 3a). The reduced thickness of the tunica media in Group-3 mice was also restored in Group-1 mice (Extended Data Fig. 6c). Ageing-and progeria-associated epigenetic marks, namely decreased histone H3 lysine 9 trimethylation (H3K9me3) and increased histone H4 lysine 20 trimethylation (H4K20me3)^20,21^, were present in Group-3 mice and were reversed in Group-1 mice (Extended Data Fig. 6d,e). In the skin, Group-3 mice showed skin atrophy, as indicated by reduced thickness of the dermis (Fig. 3b) and dermal white adipose tissue (Extended Data Fig. 6f). In contrast, Δ133p53α-expressing Group-1 mice maintained thick dermis and dermal white adipose tissue comparable to wild-type mice (Fig. 3b and Extended Data Fig. 6f). Given the alopecia and rough coat observed in HGPS patients^1^ and model mice^2^ (Extended Data Fig. 6g), respectively, we performed immunofluorescence (IF) analysis for the hair follicle stemness protein Sox9 (ref. 22). This analysis revealed diminished Sox9 expression in hair follicles of Group-3 mice, in sharp contrast to its abundant expression in those of Group-1 and wild-type mice (Fig. 3c). Group-1 mice also exhibited higher levels of Sox9 mRNA expression in the skin than Group-3 mice (Fig. 3d). In MEFs with and without progerin expression (Fig. 3e), as well as in fibroblasts derived from HGPS patients (Fig. 3f), 4-OHT-induced or lentiviral vector-driven expression of Δ133p53α increased Sox9 mRNA expression, suggesting that Δ133p53α-induced Sox9 upregulation in Group-1 mice reflects a direct regulatory effect rather than a long-term *in vivo* adaptive response.

**Fig. 3.**
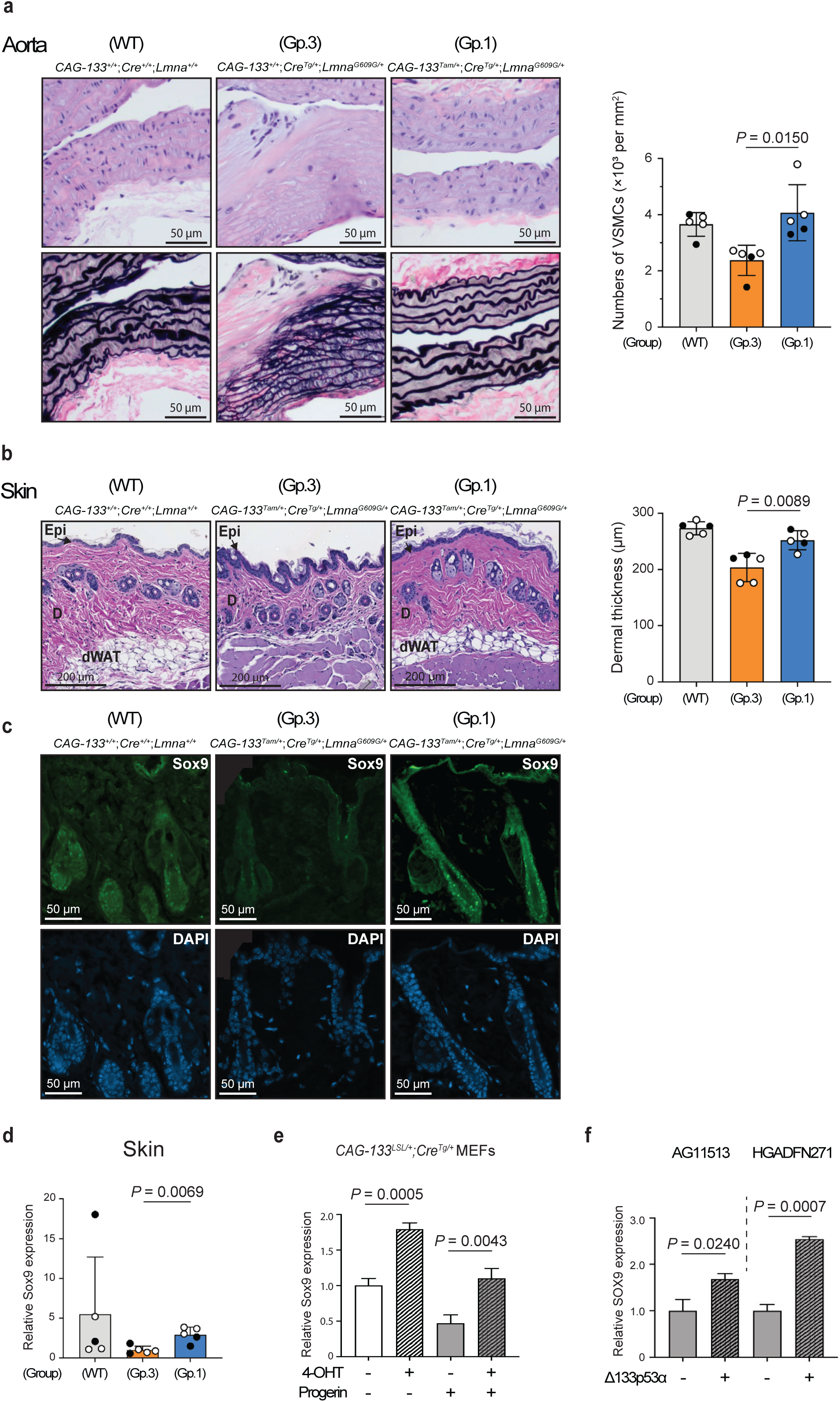
Δ133p53α abrogates the pathological changes in the aorta and skin of *Lmna^G609G/+^*mice. Tissue sections of the aorta and skin were prepared from Group-1, Group-3, and wild-type (WT) mice at 9-10 months of age (n = 5 each; open circles, females; closed circles, males). **a,** Restoration of vascular smooth muscle cells (VSMCs) in the aorta by Δ133p53α. Representative images of the serial sections stained with hematoxylin and eosin (H&E, top) and Verhoeff–Van Gieson (VVG, bottom) are shown (scale bars, 50 µm). The tunica media was identified by black-stained elastic fibers. Cell numbers in the tunica media (per mm^3^) are presented as mean ± s.d. (n = 5). **b,** Restoration of dermal thickness in the skin by Δ133p53α. Thickness of the dermis (µm) for each mouse was calculated as the average of measurements taken at five randomly selected, approximately evenly spaced sites. Scale bars, 200 µm. Data are presented as mean ± s.d. (n = 5). Epi, epidermis; D, dermis; dWAT, dermal white adipose tissue. **c,** Immunofluorescence (IF) staining of Sox9 in the hair follicles. Representative images of Sox9 (green) and DAPI (blue) are shown (scale bars, 50 µm). All five mice in each group showed highly reproducible results. **d,** Sox9 mRNA expression analyzed by qRT-PCR in the skin of Group-1, Group-3 and wild-type mice. Data are presented as mean ± s.d. (n = 5, each with technical triplicate). **e,** Sox9 mRNA expression in MEFs. *CAG-133^LSL/+^*;*Cre^Tg/+^* MEFs were with or without retroviral progerin expression, and with or without 4-OHT-induced Δ133p53α expression. **f,** SOX9 mRNA expression in HGPS patient-derived fibroblasts. Two fibroblast strains, AG11513 and HGADFN271, were transduced with Δ133p53α-expressing (+) or control (-) lentiviral vectors. Data are presented as mean ± s.d. (n = 3, technical triplicate) (**e,f**). *P* values were calculated by Welch’s *t*-test (**a,b,d,e,f**).

### Δ133p53α extends progeria mouse lifespan

Group-1 mice (*CAG-133^Tam/+^*;*Cre^Tg/+^*;*Lmna^G609G/+^*), when females and males were analyzed together, exhibited a median lifespan of 387 days, representing an approximately 11% increase (*P* = 0.0379) compared with 349 days in Group-3 mice (*CAG-133^+/+^*;*Cre^Tg/+^*;*Lmna^G609G/+^*) (Fig. 4a). A significant lifespan extension in Group 1 was also observed relative to another control group (Group 4), with median lifespans of 387 versus 358 days (*P* = 0.0312, females and males combined) (Fig. 4b). When females and males were analyzed separately, the differences did not reach statistical significance, although males still showed borderline trends (*P* = 0.0727 for Group 1 versus Group 3, and *P* = 0.0733 for Group 1 versus Group 4) (Extended Data Fig. 7a,b). Group-1 and Group-3 mice had similar body weights, both remaining underweight relative to the counterparts on the wild-type background (Extended Data Fig. 7c). Spinal kyphosis, a hallmark of ageing-associated loss of bone homeostasis^1,2^, was quantitatively assessed in Group-1 versus Group-3 mice, where the kyphosis index (inversely correlated with the severity) was significantly improved in Group-1 mice (Fig. 4c). Weekly monitoring of all Group-1 and Group-3 mice further indicated that Group-1 mice exhibited a lower incidence and later onset of spinal kyphosis compared with Group-3 mice (Fig. 4d).

**Fig. 4.**
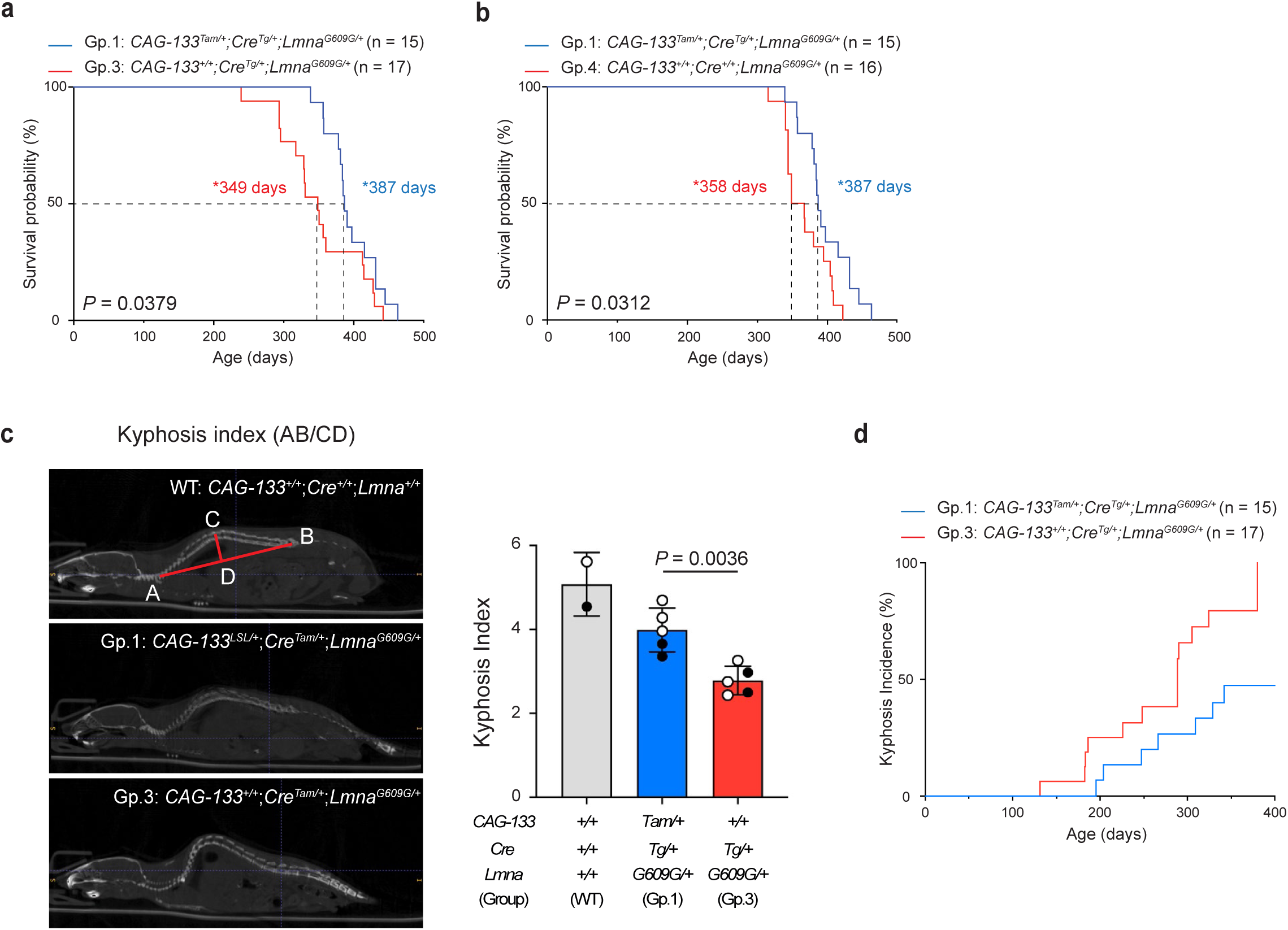
Δ133p53α extends lifespan in *Lmna^G609G/+^* mice. a,b, Kaplan-Meier survival curves comparing Group-1 mice (n = 15) with Group-3 mice (n = 17) (**a**) or Group-4 mice (n = 16) (**b**). Both female and male mice were included. Median survival days (*) and *P* values (log-rank test) are indicated. **c,** Spinal kyphosis quantitated using CT imaging. Shown on the left are representative images of a wild-type (WT) mouse, illustrating the calculation of the kyphosis index (AB divided by CD), a Group-1 mouse, and a Group-3 mouse. Quantitative comparison was performed between Group-1 and Group-3 mice (mean ± s.d. from n = 5; open circles indicate three females, and closed circles indicate two males). *P* value was determined using Welch’s *t*-test. Two wild-type mice (one female and one male) were included as references only and were not used for statistical comparison. **d,** Kyphosis incidence evaluated by gross macroscopic observation of spinal curvature. To determine the age of onset of spinal kyphosis, Group-1 (n = 15) and Group-3 (n = 17) mice were monitored weekly until 380 days of age, by which time all Group-3 mice had developed spinal kyphosis.

### Δ133p53α enhances metabolic pathways

To gain further mechanistic insight into the *in vivo* functions of Δ133p53α, we examined Group-1 versus Group-3 mice (9-10 months of age, n = 5 each) by bulk RNA-sequencing (RNA-seq) of the heart (Fig. 5a) and kidney (Fig. 5b), two organs highly vulnerable to ageing-associated changes. As expected, the p53 pathway was downregulated in both organs in Group-1 mice (Fig. 5c,d), with p53 target genes involved in cellular senescence, such as Cdkn1a (p21^Waf1/Cip1^), Btg2 and Gadd45a, as leading edge genes (Extended Data Table 2). Consistent with the repression of IL-6 by Δ133p53α (Fig. 2g,h), the pathways corresponding to increased inflammatory states, including the TNFα signaling via NFkB (Extended Data Fig. 8a,b) and the inflammatory response (Extended Data Fig. 8c), were downregulated in Group-1 mice. Of note, the oxidative phosphorylation was a highly ranked, upregulated pathway in both heart and kidney in Group-1 versus Group-3 mice (Fig. 5a,b,e,f), suggesting that Δ133p53α may contribute to enhanced mitochondrial ATP production. In the kidney, the glycolysis pathway was also highly upregulated in Group-1 mice (Fig. 5b,g), likely reflecting a hybrid metabolic state in this organ^23^. Several genes representative of the oxidative phosphorylation pathway (Ndufs6, Ndufc2, Uqcrq, Hsd17b10, and Gpx4; Extended Data Table 2) were examined by qRT-PCR and showed an overall trend toward upregulation in Group-1 mice in the heart (Fig. 5h) and kidney (Fig. 5i).

**Fig. 5.**
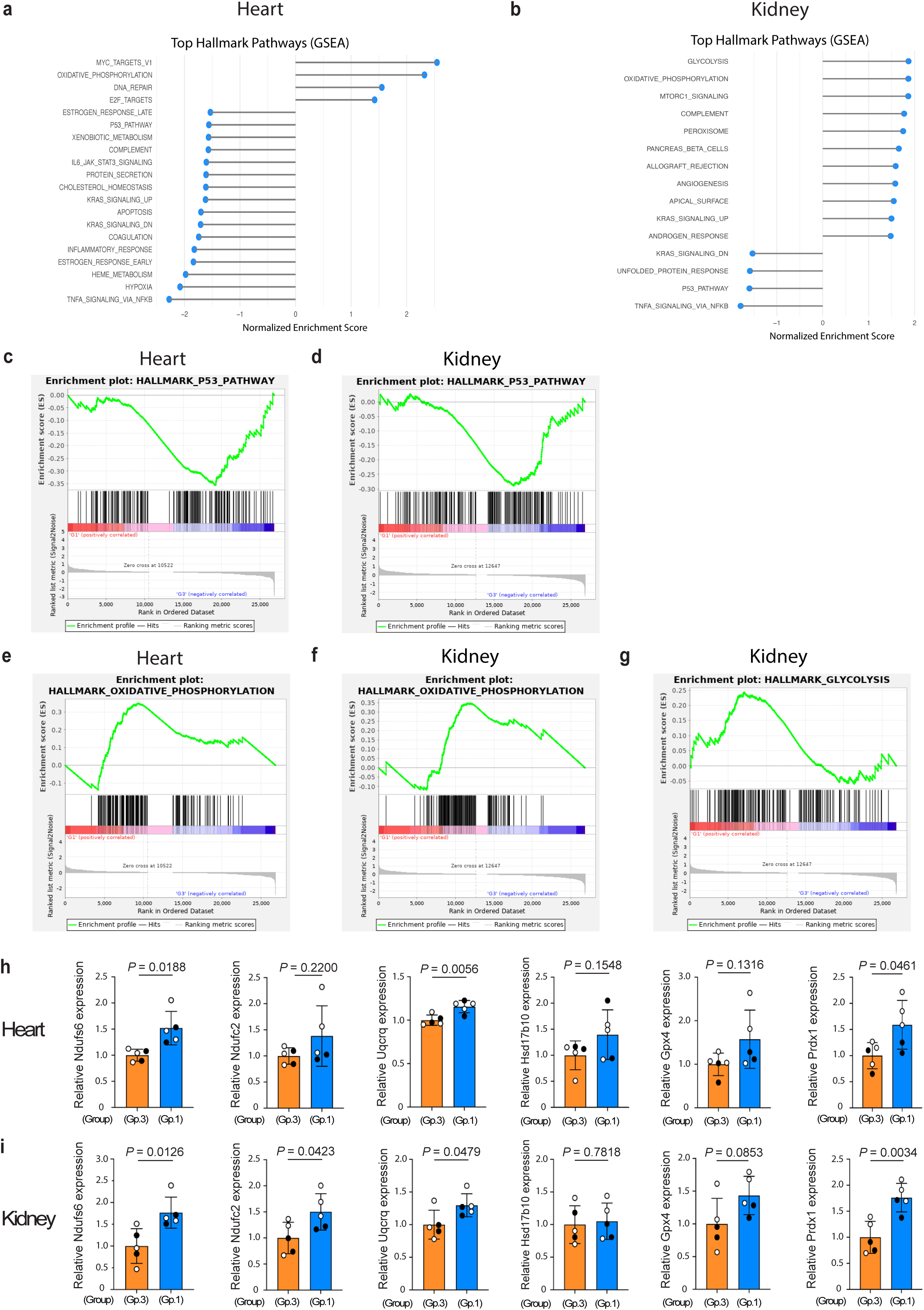
Δ133p53α enhances the gene expression profile for improved metabolism. a,b, Top Hallmark pathways identified by Gene set enrichment analysis (GSEA). The bulk RNA-seq data were obtained from the heart (**a**) and kidney (**b**) in 9-10-month-old Group-1 and Group-3 mice (n = 5 each). Pathways are ranked by normalized enrichment score. False discovery rate < 0.10. **c-g,** Enrichment plots illustrate significant downregulation of the p53 pathway (**c,d**) and upregulation of the oxidative phosphorylation (**e,f**) in both heart and kidney, as well as upregulation of the glycolysis in the kidney (**g**). Enrichment plots depict running enrichment scores (ES) and the distribution of genes within each Hallmark gene set. All leading edge genes in each pathway are listed in Extended Data Table 2. **h,i,** qRT-PCR assays of mRNA expression of genes in the oxidative phosphorylation pathway (Ndufs6, Ndufc2, Uqcrq, Hsd17b10, and Gpx4) and an antioxidant gene Prdx1 in the heart (**h**) and kidney (**i**) of 9-10-month-old Group-1 and Group-3 mice (mean ± s.d. from n = 5, each with technical triplicate; open circles, females; closed circles, males). *P* values were calculated by Welch’s *t*-test.

Among these genes, Gpx4 functions as an antioxidant enzyme. In addition, Prdx1, another antioxidant enzyme belonging to the peroxisome pathway (Extended Data Fig. 8d, Extended Data Table 2), was also upregulated in the heart (Fig. 5h) and kidney (Fig. 5i) of Group-1 mice.

### Effects of Δ133p53α in wild-type mice

To examine the effects of transgenic expression of Δ133p53α on the wild-type background, naturally aged Group-1 (*CAG-133^Tam/+^*;*Cre^Tg/+^*), Group-2 (*CAG-133^LSL/+^*;*Cre^Tg/+^*), and Group-4 (*CAG-133^+/+^*;*Cre^+/+^*, wild-type) mice were euthanized at 26-28 months of age (n = 4 each).

Although the numbers of p21^Waf1/Cip1^-, p16^Ink4a^-or γ-H2AX-positive nuclei in these naturally aged mice did not reach levels sufficient for quantitative analysis by our IHC examination, histopathological analyses of the aorta (aortic arch) and skin (dorsal back) indicated tissue-preserving effects of Δ133p53α. Compared with Group-2 and Group-4 mice, Group-1 mice showed increased numbers of VSMCs in the aortic tunica media, although no increase in thickness of the tunica media was observed (Fig. 6a). In the skin, Group-1 mice exhibited increased thickness of the dermal white adipose tissue, but not of the dermis, compared with Group-2 and Group-4 mice (Fig. 6b). Serum IL-6 levels were significantly lower in Group-1 mice when compared with the combined Group-2 and Group-4 controls, although the difference was not statistically significant when compared with either control group individually (Fig. 6c), still suggesting an anti-inflammatory effect of Δ133p53α during natural ageing.

**Fig. 6.**
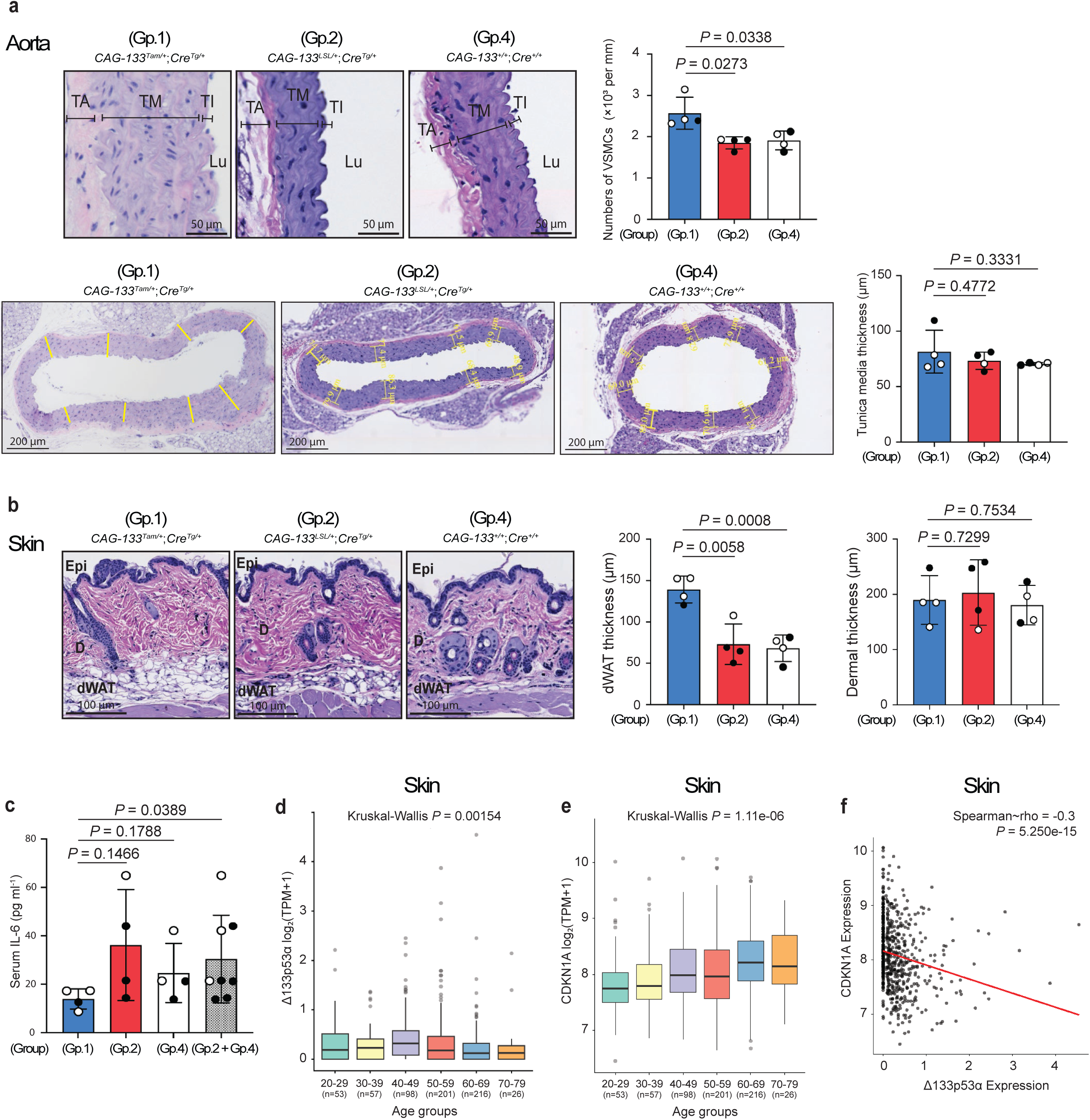
Δ133p53α effects in aged wild-type mice and Δ133p53α downregulation in aged human tissue. **a,b,** Histopathological examination of naturally aged Group-1 (*CAG-133^Tam/+^*;*Cre^Tg/+^*), Group-2 (*CAG-133^LSL/+^*;*Cre^Tg/+^*), and Group-4 (*CAG-133^+/+^*;*Cre^+/+^*, wild-type) mice at 26-28 months of age (n = 4 each; open circles, females; closed circles, males). **a,** In H&E-stained sections of the aorta, numbers of VSMCs in the tunica media (per mm^2^) were counted as in Fig. 3a. Thickness of the tunica media (µm) was measured as in Extended Data Fig. 6c. Scale bars, 50 µm (top) or 200 µm (bottom). TA, tunica adventitia; TM, tunica media; TI, tunica intima; Lu, Lumen. **b,** In H&E-stained sections of the skin, thickness of the dermal white adipose tissue as in Extended Data Fig. 6f, as well as thickness of the dermis as in Fig. 3b, was measured. Scale bars, 100 µm. Epi, epidermis; D, dermis; dWAT, dermal white adipose tissue. Data are presented as mean ± s.d. (n = 4) (**a,b**). *P* values were calculated using Welch’s *t*-test (**a,b**). **c,** Serum IL-6 concentrations (pg ml^-1^) in the same set of mice as in **a** and **b** were measured as in Fig. 2g. Data are mean ± s.d. from n = 4 (Group 1, 2, and 4) or n = 8 (Groups 2 and 4 combined). *P* values were calculated by Welch’s *t*-test. **d,e,** Analysis of human Δ133p53α (**d**) and CDKN1A (p21^WAF1/CIP1^) (**e**) expression levels in suprapubic skin tissue across age groups ranging from 20-29 to 70-79 years. Sample numbers (n) are shown in parentheses. Y-axis represents log_2_(TPM + 1). *P* values were calculated using Kruskal-Wallis test. In **d**, an outlier exceeding 10 in the 60-69 group is not shown but was included in the statistical analysis. **f,** Correlation analysis between Δ133p53α and CDKN1A expression. Spearman’s rho and *P* values are indicated (Spearman’s rank correlation test).

### Δ133p53α decreases during human ageing

The Genotype-Tissue Expression (GTEx) datasets were analyzed to assess age-associated changes in mRNA expression of Δ133p53α and CDKN1A (p21^WAF1/CIP1^) in human tissues. Due to no or low isoform-specific counts corresponding to Δ133p53α, most tissues were not suitable for statistical analysis. Nevertheless, in the aorta, the percentage of donors showing no Δ133p53α counts was higher in the >50-year groups (78%) than in the ≤ 49-year groups (64%). Notably, in suprapubic skin tissue, which allowed statistical analysis across decade-based age groups ranging from 20-29 to 70-79 years (Fig. 6d), Δ133p53α expression levels decreased significantly with increasing age. In contrast, CDKN1A expression showed a statistically significant age-dependent increase (Fig. 6e). Correlation analysis demonstrated a significant inverse relationship between Δ133p53α and CDKN1A expression (Fig. 6f). These findings suggest that Δ133p53α negatively regulates CDKN1A (p21^WAF1/CIP1^) in humans *in vivo*, consistent with such negative regulation in mice as shown in this study and in cultured human cells *in vitro* as shown previously^3,9,10^, and that endogenous regulation of Δ133p53α plays a role in natural ageing in humans.

## Discussion

This study provides evidence that Δ133p53α functions *in vivo* to counteract accelerated aging phenotypes and extend lifespan in the progeria mouse model. Transgenic expression of Δ133p53α *in vivo* not only recapitulates its cellular effects as previously observed *in vitro*, including reduced senescence (p21^Waf1/Cip1^, p16^Ink4a^, and SA-β-gal), decreased IL-6 production, and reduced DNA damage (γ-H2AX), but also mitigates the disease-characteristic pathological changes in the aorta, skin, and spine.

Genetic modifications that extend cellular replicative lifespan *in vitro* do not often result in an extension of organismal lifespan *in vivo*. For example, p53 knockout^24^ and p16^Ink4a^ knockout^25^, while extending cellular replicative lifespan *in vitro*, caused spontaneous tumorigenesis in mice early in life, leading to shortened lifespan and precluding evaluation of tumor-free natural lifespan. p21^Waf1/Cip1^-knockout mice^26^ also developed spontaneous tumors later in life, still hampering evaluation of natural lifespan. To our knowledge, the telomerase reverse transcriptase (Tert)^27^ and the pluripotent reprogramming factors (namely Oct4, Sox2, Klf4 and c-Myc, collectively termed OSKM)^28^ are two cases that resembled Δ133p53α in exhibiting both *in vitro* cellular and *in vivo* organismal lifespan-extending activities. Lentiviral expression of Tert inhibited endothelial cell senescence and extended the lifespan in a HGPS mouse model^29^. Pharmacologic activation of Tert in naturally aged mice reduced cellular senescence, inflammation and DNA damage, although the impact on lifespan remained to be determined^30^. Oct4, with or without the other reprogramming factors, also mitigated cellular senescence, inflammation and DNA damage and extended the lifespan in naturally aged mice and HGPS model mice^20,31^. Whereas continuous induction of OSKM increased tumorigenesis and mortality in HGPS model mice, CRISPR-based activation of endogenous Oct-4 or cyclic and transient induction of OSKM managed to avoid this concern of cancer-related death^20,31^. It should be noted that, in this study, continuous expression of Δ133p53α after induction at 5-6 weeks of age did not increase cancer-related death in heterozygous *Lmna^G609G^* HGPS mice (Fig. 4a,b and Extended Data Table 1). Over a longer term during natural ageing on the wild-type background, continuous expression of Δ133p53α after 8-10 weeks of age also did not increase lethal spontaneous tumors (Extended Data Table 1). This non-tumorigenic nature *in vivo* of continuous Δ133p53α expression directly validates our *in vitro* findings that Δ133p53α did not cause malignant transformation or genomic instability in human cells^3,9,10^.

We speculate that the fundamental nature of Δ133p53α-mediated lifespan extension is rooted in its inhibitory effect on p53-mediated senescence^3,9,10^, which is represented by reduced expression of p21^Waf1/Cip1^ (Fig. 1a) and is associated with cellular stemness^10^. As mentioned above, robust and unregulated inhibition of p53 or p21^Waf1/Cip1^, such as through genetic knockout, leads to increased tumorigenesis and associated mortality during long-term natural ageing^24,26^, masking any potential lifespan-extending effects in the absence of tumors. Under experimental settings using shorter-lived models and/or significant but regulated inhibition of p21^Waf1/Cip1^ as adopted in this study, the potential of extending lifespan is poised to be revealed. For example, in *Zmpste24*^-/-^ progeroid mice with a median lifespan of approximately 100 days, p53 knockout resulted in significant extension of lifespan with reduced expression of p21^Waf1/Cip1^ (ref. 32).

MicroRNA-302b, which has been reported to negatively regulate p53 (ref. 33), extended lifespan in naturally aged wild-type mice, with reduced p21^Waf1/Cip1^ expression and without an increase in tumorigenesis^34^. Based on these previous studies and our own findings in this study, we propose that an appropriate level and/or repertoire of p53 inhibition represents a physiologically non-disruptive approach for lifespan extension with no or minimal risk of tumorigenesis.

This study also suggests that Δ133p53α influences other factors and processes in favor of mitigating ageing-related pathologies and extending lifespan. Decreased serum IL-6 levels (Fig. 2g) indicate reduced levels of systemic inflammation, which also accompanied lifespan extension in HGPS model mice treated with an anti-IL-6 receptor neutralizing antibody^35^ and in wild-type mice with suppression of the insulin-IGF-mTORC-Ras network^36^. Enhanced oxidative phosphorylation in the heart and kidney (Fig. 5), which was also observed in Δ133p53α-armored chimeric antigen receptor T cells^11^, indicates improved mitochondrial metabolism^37^. In the kidney, enhanced glycolysis was also observed (Fig. 5g), possibly reflecting the difference in energy production between the renal cortex (oxidative phosphorylation) and medulla (glycolysis)^23^. The antioxidant enzymes Gpx4 and Prdx1 were upregulated by Δ133p53α (Fig. 5h.i), likely contributing to protective responses against HGPS-associated oxidative stress^38^. Aging-associated histone modifications, such as decreased H3K9me3 and increased H4K20me3 (ref. 20), were reversed by Δ133p53α expression in the aorta (Extended Data Fig. 6d,e), suggesting that Δ133p53α may also influence the epigenetic landscape of ageing. Sox9, a chromatin pioneer factor for hair follicle stem cells in the skin^22^, was induced by Δ133p53α (Fig. 3c,d), potentially promoting skin homeostasis and delaying the appearance of rough hair. Sox9 is reported to be directly downregulated by p53 (ref. 39), consistent with its upregulation by Δ133p53α in this study (Fig. 3e,f). Based on all our findings, we propose that Δ133p53α promotes multiple aspects of youth and longevity not only by directly antagonizing p53-mediated senescence but also through a broad spectrum of mechanisms involving inflammation, metabolism, antioxidant defense, epigenetics, and tissue stemness.

Our findings in this study, together with our previous studies^3,9,10,12,15^, support a therapeutic strategy utilizing Δ133p53α for functionally reprogramming otherwise senescent cells and thereby promoting organismal health^40,41^. Given that transgenic expression of Δ133p53α in wild-type mice also exhibited some beneficial effects similar to those observed in *Lmna^G609G/+^*mice (Fig. 6a-c) and that Δ133p53α levels in human skin tissue decrease with increasing age (Fig. 6d), such a Δ133p53α-based reprogramming strategy may potentially be applicable not only to progeria patients but also to the general human population. Senolytic drugs, which are capable of eliminating senescent cells, have shown protective effects against bone and muscle degeneration in progeria model mice^42,43^ and are currently being tested in clinical trials for senescence-associated diseases such as neurodegenerative diseases, idiopathic pulmonary fibrosis, and osteoarthritis^44^. We expect that Δ133p53α-based and other cellular reprogramming strategies cooperate with senolytic strategies to complementarily target heterogenous populations of senescent cells^40,41^. When considering combinatorial therapies for HGPS, it should be noted that transgenic expression of Δ133p53α did not significantly reduce progerin expression levels in most of the tissues examined in *Lmna^G609G/+^*mice (Fig. 1 and Extended Data Fig. 4c). This is in contrast to reduced progerin levels observed with other interventions in HGPS model mice, including Oct-4 activation^31^, IL-6 neutralization^35^, and Tert expression^29^. The progerin-independent anti-HGPS action of Δ133p53α prompts its combination with currently available progerin-targeting therapies^5^, potentially leading to an effective combinatorial therapy based on mechanistically distinct strategies.

The Δ133p53α-transgenic mice, newly generated and utilized in this study for ubiquitous, postnatal induction of Δ133p53α, will be applicable to a wide range of mouse disease models (for example, neurodegenerative disease models and idiopathic pulmonary fibrosis models), in which ubiquitous or tissue-specific induction of Δ133p53α may exert therapeutic effects^41^.

However, our experimental settings in this study may also highlight limitations associated with this mouse model. Considering the 3-4 weeks of weaning periods for *Lmna^G609G/+^* and *Lmna^G609G/G609G^*pups^2^ and the additional time needed for genotyping, the five-day consecutive i.p. injections of tamoxifen in this study were performed at 5-6 weeks of age, which might not have been early enough to capture the full effects of Δ133p53α on progeria-related phenotypes. In support of this notion, Δ133p53α-induced lifespan extension did not reach statistical significance in shorter-lived homozygous *Lmna^G609G/G609G^*mice, although Δ133p53α still significantly inhibited loss of body weight in these mice (Extended Data Fig. 9). These findings underscore the need to further optimize Δ133p53α induction protocols in transgenic mouse models, as well as to develop non-genetic, pharmacological methods for Δ133p53α activation. Two compounds recently identified to increase Δ133p53α protein levels in human cells^45,46^, together with this present study demonstrating the *in vivo* activities of Δ133p53α, should facilitate the clinical translation of Δ133p53α.

## Methods

### Mice

All animal studies have been approved by the Animal Care and Use Committee (ACUC) at National Cancer Institute (NCI, Frederick, MD) under the protocols ASP 25-264, ASP 24-466 and ASP 21-0212. All animal procedures, including housing and environmental enrichment, breeding, genotyping, recognition and alleviation of pain and distress, collection of tissues and blood, humane endpoints, and euthanasia, followed the ACUC guidelines (https://ncifrederick.cancer.gov/Lasp/ACUC/Frederick/GuidelinesFnl). The HGPS model mice carrying the *Lmna^G609G^* allele on the C57BL/6J background^4^ were a gift from Dr. Carlos López-Otín (University of Oviedo) through Dr. Francis Collins (National Human Genome Research Institute). The UBC-Cre-ERT2 strain on the C57BL/6J background^47^, which ubiquitously expresses a tamoxifen-activated Cre recombinase, was obtained from Jackson Laboratory (stock no. 007001).

To achieve an inducible, transgenic allele of Δ133p53α on the C57BL/6J background, we first constructed the targeting vector for knocking-in at the *ROSA26* locus (Extended Data Fig. 1a). This targeting vector was co-injected with Cas9 mRNA and a gRNA targeting the *ROSA26* sequence (CTCCAGTCTTTCTAGAAGATGGG) into fertilized eggs, followed by identification of F0 founder mice by PCR-genotyping and Sanger sequencing (the sequences of PCR and sequencing primers are available upon request). Two F0 founders were bred to wild-type mice to confirm germline transmission and generate F1 mice. All the procedures through obtaining F1 mice were conducted at Cyagen Biosciences under the project number ROSAM-200325-ADJ-01.

The resulting transgenic allele of Δ133p53α (CAG-LSL-Δ133p53α) was maintained by breeding the heterozygotes (termed *CAG-133^LSL/+^*) with wild-type mice. *CAG-133^LSL/+^* mice were crossed with heterozygous UBC-Cre-ERT2 mice (termed *Cre^Tg/+^*) to obtain *CAG-133^LSL/+^*;*Cre^Tg/+^*mice. *CAG-133^LSL/+^*;*Cre^Tg/+^* mice were bred with wild-type mice (*CAG-133^+/+^*;*Cre^+/+^*) to maintain this crossbred strain and to obtain the experimental and control groups of naturally aged mice. For experiments on the heterozygous *Lmna^G609G^* background, heterozygous *Lmna^G609G^*mice (termed *Lmna^G609G/+^*) were crossed with *CAG-133^LSL/+^*;*Cre^Tg/+^*mice to obtain *CAG-133^LSL/+^*;*Cre^Tg/+^*;*Lmna^G609G/+^*mice as well as the other two genotypes used: *CAG-133^+/+^*;*Cre^Tg/+^*;*Lmna^G609G/+^*and *CAG-133^+/+^*;*Cre^+/+^*;*Lmna^G609G/+^*. For experiments on the homozygous *Lmna^G609G^* background, *CAG-133^LSL/+^*;*Cre^Tg/+^*;*Lmna^G609G/+^*mice were bred with *Lmna^G609G/+^* mice to obtain the three genotypes used: *CAG-133^LSL/+^*;*Cre^Tg/+^*;*Lmna^G609G/G609G^*, *CAG-133^+/+^*;*Cre^Tg/+^*;*Lmna^G609G/G609G^* and *CAG-133^+/+^*;*Cre^+/+^*;*Lmna^G609G/G609G^*. In all our breeding strategies, both CAG-LSL-Δ133p53α and UBC-Cre-ERT2 alleles were heterozygous whenever present, which were detected by the following genotyping PCR primers: TTTTGCCAACTGGCCAAGACCTGC and GTCTGAGTCAGGCCCTTCTGTCTTG for CAG-LSL-Δ133p53α; and GACGTCACCCGTTCTGTTG and AGGCAAATTTTGGTGTACGG for UBC-Cre-ERT2. *Lmna^+/+^*, *Lmna^G609G/+^* and *Lmna^G609G/G609G^*genotypes were distinguished by PCR using the primers AAGGGGCTGGGAGGACAGAG and AGCATGCAATAGGGTGGAAGGA, which generates 100-bp (*^+^*, wild-type) and 240-bp (*^G609G^*, mutant) products.

Intraperitoneal (i.p.) injection of tamoxifen was performed to induce Cre recombination, as previously established^48^. The i.p. injection was performed on five consecutive days to maximize the efficiency of Cre recombination. In heterozygous and homozygous *Lmna^G609G^* studies, the i.p. injection of tamoxifen into *CAG-133^LSL/+^*;*Cre^Tg/+^*;*Lmna^G609G/+^*and *CAG-133^LSL/+^*;*Cre^Tg/+^*;*Lmna^G609G/G609G^*mice was performed at 5-6 weeks of age (renamed as *CAG-133^Tam/+^*;*Cre^Tg/+^*;*Lmna^G609G/+^* and *CAG-133^Tam/+^*;*Cre^Tg/+^*;*Lmna^G609G/G609G^* after tamoxifen injection, respectively, and called Group 1 in each study). Some of the *CAG-133^LSL/+^*;*Cre^Tg/+^*;*Lmna^G609G/+^*mice (heterozygous study) did not receive tamoxifen injection (called Group 2, representing no-tamoxifen controls). All *CAG-133^+/+^*;*Cre^Tg/+^*;*Lmna^G609G/+^*(heterozygous study) and *CAG-133^+/+^*;*Cre^Tg/+^*;*Lmna^G609G/G609G^*mice (homozygous study) were injected with tamoxifen at 5-6 weeks of age (called Group 3 in each study, representing tamoxifen-treated, Cre-activated controls lacking the Δ133p53α transgene). All *CAG-133^+/+^*;*Cre^+/+^*;*Lmna^G609G/+^*(heterozygous study) and *CAG-133^+/+^*;*Cre^+/+^*;*Lmna^G609G/G609G^*mice (homozygous study) were left non-injected (called Group 4 in each study, representing no-transgene controls). In a study on the wild-type background, the i.p. injection of tamoxifen into *CAG-133^LSL/+^*;*Cre^Tg/+^*mice was performed at 8-10 weeks of age (renamed as *CAG-133^Tam/+^*;*Cre^Tg/+^*after tamoxifen injection and called Group 1). Some *CAG-133^LSL/+^*;*Cre^Tg/+^*mice and all *CAG-133^+/+^*;*Cre^+/+^* mice were left non-injected (the former called Group 2, representing no-tamoxifen controls, and the latter called Group 4, representing no-transgene controls).

To summarize above, Group 1 represents the experimental group with transgenic Δ133p53α induced, Group 2 is a control group with the same genotype as Group 1 but without tamoxifen injection, Group 3 is a control group lacking the Δ133p53α transgene but carrying Cre recombinase, which was activated by tamoxifen injection, and Group 4 is a control group without either transgene. In this manuscript, all mouse experiments included either Group 2 or Group 3 as the optimal control; in most cases, Group 4 and/or wild-type mice were also included as additional controls. Both male and female mice were used, with sex specified where relevant.

### Collection of samples and examination of lifespan in mice

Scheduled euthanasia of age-matched mice from the experimental group (Group 1) and two or three control groups (Groups 2, 3 and/or 4) was carried out to collect serum samples, prepared from blood obtained via post-mortem cardiac puncture, as well as tissue samples, which were snap-frozen or fixed in 10% neutral buffered formalin (NBF). For lifespan analysis, 10-17 mice assigned per group (exact numbers of mice are mentioned in each experiment) were monitored daily for general health conditions. Body weight was measured weekly (in heterozygous and homozygous *Lmna^G609G^* mice) or every other week (in natural aged mice), along with monitoring for hair loss and visible or palpable tumors. Humane endpoints for individual mice followed the ACUC 10.00 guidelines (https://ncifrederick.cancer.gov/Lasp/ACUC/Frederick/GuidelinesFnl), including 20% loss of body weight, difficulty reaching food, difficulty breathing, and morbidity. Mice exhibiting any of these conditions were euthanized and necropsied, and when needed, their serum and tissue samples were collected as described above for scheduled euthanasia. The day on which mice were found dead or euthanized due to humane endpoints was defined as the end of their lifespan. Lifespan was analyzed for the experimental group (Group 1) versus a single control group (Group 3 or 4). Kaplan-Meier survival curves and log-rank tests were performed using GraphPad Prism v.10.6.1. All mice assigned to the lifespan analysis and scheduled euthanasia are listed with their date of birth, date of death or euthanasia, specific abnormalities or endpoint criteria, and lifespan in Extended Data Table 1.

### Western blot analysis

Snap-frozen tissues were homogenized in the lysis buffer (Thermo Fisher Scientific, 89900) with the protease/phosphatase inhibitor cocktail (Thermo Fisher Scientific, 78440) using PowerLyzer 24 (QIAGEN) with ceramic beads (QIAGEN, 13114-50), followed by centrifugation at 12,000*g* for 20 min. Cultured cells were suspended in the same lysis buffer and centrifuged. Gel electrophoresis, transfer to PVDF membranes, membrane blocking, incubation with primary antibodies and horse radish peroxidase (HRP)-conjugated secondary antibodies, and membrane washing were carried out as previously described^10,15^. The SuperSignal West Dura Extended Duration Substrate (Thermo Fisher Scientific, 34076) was used as an HRP substrate for chemiluminescence, which was detected on the ChemiDoc Imaging System (Bio-Rad).

Quantitative image analysis was performed using the Image Lab software ver. 6.1 (Bio-Rad). Primary antibodies used were as follows: anti-p53 antibodies SAPU (sheep polyclonal)^49^ and DO-11 (mouse monoclonal; Bio-Rad, MCA1704)^50^; anti-p21^Waf1/Cip1^ (mouse monoclonal; Santa Cruz Biotechnology, sc-6246, clone F-5); anti-lamin A/C (mouse monoclonal; Santa Cruz Biotechnology, sc-376248); anti-GAPDH (mouse monoclonal; Santa Cruz Biotechnology, sc-166574); and anti-β-actin (mouse monoclonal; Thermo Fisher Scientific, MA1-91399). HRP-conjugated secondary antibodies used were goat anti-mouse IgG (H+L) (Thermo Fisher Scientific, 32430), goat anti-rabbit IgG (H+L) (Thermo Fisher Scientific, 32460), and donkey anti-sheep IgG (H+L) (Jackson ImmunoResearch, 713-035-147).

### RNA isolation and quantitative RT-PCR (qRT-PCR)

Total RNA samples were isolated using the RNeasy Plus Micro Kit (QIAGEN, 74034). Snap-frozen tissues were homogenized, as above, in the RLT Plus buffer (supplemented with β-mercaptoethanol) provided in the kit. Cultured cells were suspended in the same RLT Plus buffer. Subsequently, both tissue and cell samples were processed according to the manufacturer’s instructions. RNA quality was checked on the Agilent TapeStation at the CCR Genomics Core, National Cancer Institute (NCI). Reverse transcription for cDNA synthesis was performed using the High-Capacity cDNA Reverse Transcription Kit (Thermo Fisher Scientific, 4368814). qRT-PCR assays were performed on the 7500 Real-Time PCR system (Applied Biosystems) using the Taqman Gene Expression Master Mix (Thermo Fisher Scientific, 4369016) and the following primers/probe sets (Thermo Fisher Scientific): Cdkn1a (Mm00432448_m1), Il6 (Mm00446190_m1), Ndufs6 (Mm02529639_u1), Ndufc2 (Mm02526996_g1), Uqcrq (Mm03052986_s1), Hsd17b10 (Mm00840109_m1), Gpx4 (Mm04411498_m1), Prdx1 (Mm01621996_s1), Sox9 (Mm00448840_m1), and SOX9 (Hs00165814_m1), as well as Gapdh (Mm99999915_g1) and GAPDH (Hs02758991_g1) as internal controls. Quantitative data analysis was performed using the ΔΔCt method (https://assets.thermofisher.com/TFS-Assets/LSG/manuals/cms_042380.pdf). All samples were analyzed in technical triplicate, yielding highly reproducible Ct values with standard deviations of less than 0.15 in all assays.

### Histopathological examination

Aorta, skin and muscle tissue samples fixed in 10% NBF were processed and embedded in paraffin to prepare formalin-fixed paraffin-embedded (FFPE) blocks. Sections were cut from the FFPE blocks at a thickness of 5 µm, followed by staining with hematoxylin and eosin (H&E), at the Molecular Histopathology Laboratory, NCI-Frederick. The H&E-stained slides were assessed in a blinded manner by a board-certified veterinary pathologist at the Laboratory Animal Sciences Program, NCI-Frederick.

### Immunohistochemical (IHC) and immunofluorescence (IF) staining

FFPE slides were deparaffinized and rehydrated, followed by heat-induced antigen retrieval in Antigen Unmasking Solution (Vector Laboratories, H-3300). For IHC, endogenous peroxidase activity was blocked with BLOXALL® Endogenous Enzyme Blocking Solution (Vector Laboratories, SP-6000). Primary antibodies used were: anti-p21^Waf1/Cip1^ (Santa Cruz Biotechnology, sc-6246; 1:100), anti-p16^Ink4a^ (Santa Cruz Biotechnology, sc-1661; 1:100), anti-γ-H2AX (MilliporeSigma, JBW301; 1:200), anti-Sox9 (MilliporeSigma, AB5535; 1:200), anti-histone H3K9me3 (Abcam, ab8898; 1:500), and anti-histone H4K20me3 (Abcam, ab9052; 1:200). For IHC, immunodetection was performed using the M.O.M. Immunodetection Kit (Vector Laboratories, BMK-2202) for mouse primary antibodies, or the VECTASTAIN Elite ABC Kit (Vector Laboratories, PK-6101) for rabbit primary antibodies, followed by visualization using ImmPACT DAB Substrate Kit (Vector Laboratories, SK-4105), hematoxylin counterstaining, dehydration, clearing, and mounting. Secondary antibodies for IF were: donkey anti-rabbit IgG (H+L) highly cross-adsorbed secondary antibody, Alexa Fluor 594 (Thermo Fisher Scientific, A-21207) or donkey anti-mouse IgG (H+L) highly cross-adsorbed secondary antibody, Alexa Fluor 488 (Thermo Fisher Scientific, A-21202). The slides were mounted with ProLong Gold Antifade Mountant with DAPI (4′,6-diamidino-2-phenylindole) (Thermo Fisher Scientific, P36931) and then analyzed using the Zeiss LSM 780 confocal microscope with the Zeiss ZEN software at the CCR Microscopy Core, NCI. Quantitative image analysis for both IHC and IF was performed using HALO software (Indica Labs), with nuclear or area-based quantification modules selected as appropriate for each marker.

### Verhoeff-Van Gieson staining of aorta sections

The FFPE slides prepared from aorta were deparaffinized and rehydrated. Verhoeff–Van Gieson (VVG) staining was performed at the Molecular Histopathology Laboratory, NCI-Frederick, as previously described^51^. In brief, the slides were stained for 15 min in Verhoeff’s elastic stain solution (2.5% hematoxylin, 2.5% ferric chloride, 25% ethanol, 5 mg ml^-1^ iodine, and 10 mg ml^-1^ potassium iodine), followed by differentiation in 2% ferric chloride until elastic fibers turned black with the gray background. The slides were then treated with 5% sodium thiosulfate for 1 min, rinsed in water, and counterstained for 1 min in Van Gieson solution (0.05% acid fuchsin in saturated aqueous picric acid), which distinguished collagen fibers (red), elastic fibers and nuclei (black), and other tissue structures (yellow). Following staining, the slides were dehydrated through graded ethanol, cleared in xylene, and cover-slipped. Stained slides were examined using a bright-field light microscope.

### Senescence-associated **β**-galactosidase (SA-**β**-gal) staining

**S**pleen tissues were embedded in optimal cutting temperature compound (OCT) and then snap-frozen. The OCT-embedded, snap-frozen tissues were cryo-sectioned at a thickness of 20 µm using the Cryo-Jane system (Leica Biosystems). The slides were fixed in 2% formaldehyde plus 0.2% glutaraldehyde, and stained with 1 mg ml^-1^ 5-bromo-4-chloro-3-indolyl-β-D-galactopyranoside (X-gal) at pH 6.0, using the Senescence β-Galactosidase Staining Kit (Cell Signaling Technology, 9860). The slides were then counterstained with 0.1% neutral red solution for 1 min, dehydrated in 100% ethanol, washed with xylene, and cover-slipped. Image acquisition and quantification were performed with the HALO software Area Quantification module (Indica Labs). Cells in cell culture were fixed and stained using the same staining kit, followed by cell counting under microscopic observation.

### Enzyme-linked immunosorbent assay (ELISA)

Serum samples (50 µl) were quantitated using the Mouse IL-6 Quantikine ELISA Kit (R&D Systems, M6000B). The experiment was carried out in 96-well plate format, where the absorbance was measured at 450 nm using the SpectraMax ABS Plus microplate reader (Molecular Devices). Serial dilutions of recombinant mouse IL-6 were used to generate a standard curve ranging from 0 to 250 pg ml^-1^. Each sample was analyzed in technical duplicates, which gave highly reproducible values.

### Kyphosis index

Severity of spinal kyphosis was evaluated by calculating the kyphosis index, as previously described^52^. In brief, mice were anesthetized with isoflurane and imaged by computed tomography (CT) in lateral projection using the Mediso nanoScan SPEC/PET/CT system at the Small Animal Imaging Program, NCI-Frederick. A straight-line distance between the caudal margin of the last cervical vertebra and the caudal margin of the sixth lumbar vertebra (line AB) and a perpendicular distance from this line to the dorsal edge of the vertebra at the point of maximum curvature (line CD) were measured. The kyphosis index (AB divided by CD) decreases with increasing severity. The presence or absence of spinal kyphosis was also visually monitored weekly to determine the onset of the disease^53^.

### Bulk RNA-sequencing (RNA-seq)

Total RNA samples were treated with DNase I (Thermo Fisher Scientific, 18068015) and checked for quality using the Agilent TapeStation. The RNA integrity numbers (RIN) were 8.0 or higher. RNA-seq libraries were prepared using Illumina Stranded mRNA Prep and sequenced on an Illumina NovaSeq X Plus platform using paired-end sequencing (2 × 100 bp) at the Frederick Sequencing and Genomics Core, NCI-Frederick. Sequencing reads were trimmed for adapters and low-quality bases using Cutadapt and aligned to the mouse reference genome (mm10; GENCODE M2 annotation) using STAR. Mapping statistics were calculated using Picard. Gene-level expression quantification was performed using STAR/RSEM to generate raw and normalized count matrices. Differential gene expression analysis was conducted using DESeq2, with significance defined by P < 0.05 and an absolute log_2_ fold change > 1. Pathway and functional enrichment analyses were performed using Ingenuity Pathway Analysis (IPA; QIAGEN) and Gene Set Enrichment Analysis (GSEA), focusing on Hallmark gene sets.

### Preparation of mouse embryonic fibroblasts (MEFs)

Mating cages were set up with *CAG-133^LSL/+^*;*Cre^Tg/+^*male and wild-type (*CAG-133^+/+^*;*Cre^+/+^*) female mice. Females with copulation plugs were transferred to a new cage and the embryos were recorded as embryonic day 0.5 (E0.5). On E14.5, those females were anesthetized by isoflurane inhalation, then euthanized by cervical dislocation. Each fetus was transferred to a 10-cm dish, washed with PBS, minced into fine pieces (approximately 1 mm). Minced pieces were transferred to a 15-ml tube, suspended in 2 ml of ice-cold 0.25% trypsin, and incubated overnight at 4 °C. After adding 2 ml of cell culture medium (Dulbecco’s modified Eagle medium containing 10% FBS, 50 U ml^-1^ penicillin and 50 μg ml^-1^ streptomycin), pipetting up and down, and centrifuging at 300*g* for 5 min (repeated twice), cells were resuspended in fresh cell culture medium on a 10-cm dish or in a T-25 flask, and maintained in a humidified incubator with 5% CO_2_ at 37 C°. *CAG-133^LSL/+^*;*Cre^Tg/+^*MEFs (for Δ133p53α induction) and *CAG-133^+/+^*;*Cre^+/+^*MEFs (wild-type control) were used. At each passage, cells were expanded at a split ratio of 1:3.

Cumulative population doubling levels (PDL) achieved between passages were calculated as log_2_ (number of cells obtained/number of cells seeded), as previously performed^9^.

### Human fibroblasts derived from patients with HGPS

AG11513 was obtained from Coriell Institute for Medical Research (https://catalog.coriell.org/). HGADFN271 was obtained from Progeria Research Foundation (https://www.progeriaresearch.org/).

### Retroviral and lentiviral vectors and transduction

The retroviral vector for expression of progerin (pBABE-puro-GFP-progerin) was obtained from Addgene (Plasmid #17663). Retroviral packaging and transduction into *CAG-133^LSL/+^*;*Cre^Tg/+^*MEFs were performed as previously described^9^. After two days of puromycin selection (2 µg ml^-1^), the cells were split into two sets: one set treated for 72 h with 1 µM 4-OHT; and the other set treated with DMSO alone. After an additional two days of culture, these MEFs were used for qRT-PCR analyses.

The lentiviral vector for expression of Δ133p53α (pLOC-Δ133p53α) and its control vector (pLOC-RFP, Open Biosystems) were previously described^15,54^. Lentiviral packaging and transduction into AG11513 and HGADFN271 fibroblasts were performed as previously described^15,54^. After four days in the presence of blasticidin (5 µg ml^-1^), the cells were used for qRT-PCR analyses.

### Analysis of human gene expression datasets

We analyzed postmortem RNA-seq data generated by the Genotype-Tissue Expression (GTEx) consortium (v8 release) from 948 donors aged 20–79 years. Only samples that passed GTEx quality control criteria and were annotated as non-pathological were included. Sequencing reads were aligned to the GRCh38 human reference genome, and transcript abundances were estimated using RSEM. Expression of Δ133p53α was queried based on its annotated transcription start site within intron 4 of the *TP53* locus and extracted using the transcript identifier ENST00000504937.5 (GENCODE v26). Expression of *CDKN1A* (ENSG00000124762.14) was assessed at the gene level using normalized transcripts per million (TPM). TPM values were log-transformed [log_2_(TPM + 1)] prior to downstream analyses to reduce skewness and heteroscedasticity. Donors were stratified into decade-based age groups (20–29, 30–39, 40–49, 50–59, 60–69, and 70-79 years). Age-associated changes in Δ133p53α and *CDKN1A* expression were evaluated using linear regression models with age as the primary independent variable.

Statistical comparisons across age groups within each tissue were performed using the Kruskal–Wallis test. Associations between Δ133p53α and *CDKN1A* expression were assessed using Spearman rank correlation analysis. All analyses were conducted in R (version 4.5), and visualization of age-dependent expression trends was performed using ggplot2.

## Statistical analyses

Statistical analyses were performed using GraphPad Prism (v.10.6.1) and R (v.4.5). All tests were two tailed. Welch’s *t*-test was performed to quantitatively analyze data obtained in western blot, qRT-PCR, IHC, ELISA, histopathology, kyphosis, and tissue SA-β-gal assays. Kaplan-Meier survival curves and log-rank tests were conducted for lifespan analyses. Body weights were analyzed using a mixed-effects model. One-way ANOVA with Tukey’s multiple comparison test was used for SA-β-gal assays in cultured cells. Nested one-way ANOVA was used to analyze γ-H2AX data in cultured cells. Differential expression analysis of RNA-seq data was performed using DESeq2. GTEx human expression data in different age groups were analyzed using Kruskal-Wallis test. Correlation analyses were performed using Spearman’s rank correlation test. Error bars represent mean ± s.d. *P* < 0.05 was considered statistically significant. Figure legends and/or figures include the statistical methods used and the exact *P*-values (unless less than 0.0001).

## Reporting summary

Further information on research design is available in the Nature Portfolio Reporting Summary linked to this article.

## Data availability

The RNA-seq data analyzed in this study have been deposited in the NCBI Gene Expression Omnibus (GEO) and are accessible through GSE315211. The CAG-LSL-Δ133p53α strain generated in this study has been deposited in the Mutant Mouse Resource & Research Centers (ID 075789-UNC). The pLOC-Δ133p53α plasmid has been deposited in the Addgene (#241916).

## Acknowledgements

We thank Jessica Ebersole, Eleazar Vega-Valle, Morgan Whipp, Jacqueline Clem and Christine Perella for animal care and support. We also thank the following NCI-Frederick Laboratory Services for their technical assistance and advice: Molecular Histopathology Laboratory (Stephanie Smith, Elijah Edmondson, Baktiar Karim and other staff); Small Animal Imaging Program (Joseph Kalen); Animal Diagnostic Laboratory (Mei-Chen Tseng, Sue Carney and other staff); and Illumina Sequencing Facility (Bao Tran, Jyoti Shetty, and Yongmei Zhao). We also appreciate CCR Genomics Core and CCR Microscopy Core for technical assistance. Deepak Joshi (Taconic Biosciences) and Mosen Chen (Cyagen Biosciences) contributed to the planning of mouse strain generation. A sheep polyclonal antibody SAPU was a gift from Jean-Christophe Bourdon. pBABE-puro-GFP-progerin was a gift from Tom Misteli through Addgene. CAG-LSL-Δ133p53α mice are maintained at the Mutant Mouse Resource and Research Center U42OD010924 for distribution. This research was supported by the Intramural Research Program of the National Institutes of Health (NIH) (ZIA BC 011496). The contributions of the NIH authors were made as part of their official duties as NIH federal employees, are following agency policy requirements, and are considered Works of the United States Government. However, the findings and conclusions presented in this paper are those of the authors and do not necessarily reflect the views of the NIH or the U.S. Department of Health and Human Services. L.Y. was supported by the JSPS Research Fellowship for Japanese Biomedical and Behavioral Researchers at NIH.

## Author contributions

I.H, N.v.M. and C.C.H. conceived the project. I.H. generated the transgenic mouse strain. L.Y. and I.H. designed and conducted all the laboratory and animal experiments. I.H. and N.v.M. accumulated knowledge and generated materials essential to the project. L.Y. and H.L. analyzed RNA-seq data and GTEx database. L.Y. and I.H. analyzed all the remaining data in the manuscript. C.C.H. acquired funding. I.H. wrote the manuscript. All authors reviewed the manuscript and approved the final version of the manuscript.

## Additional Information

**Supplementary information** is available for this paper.

**Correspondence and requests for materials** should be addressed to Izumi Horikawa.

