## Supplementary Figures 1-9 for "Senescence-inhibitory Δ133p53α counteracts accelerated ageing and mortality"

**a**

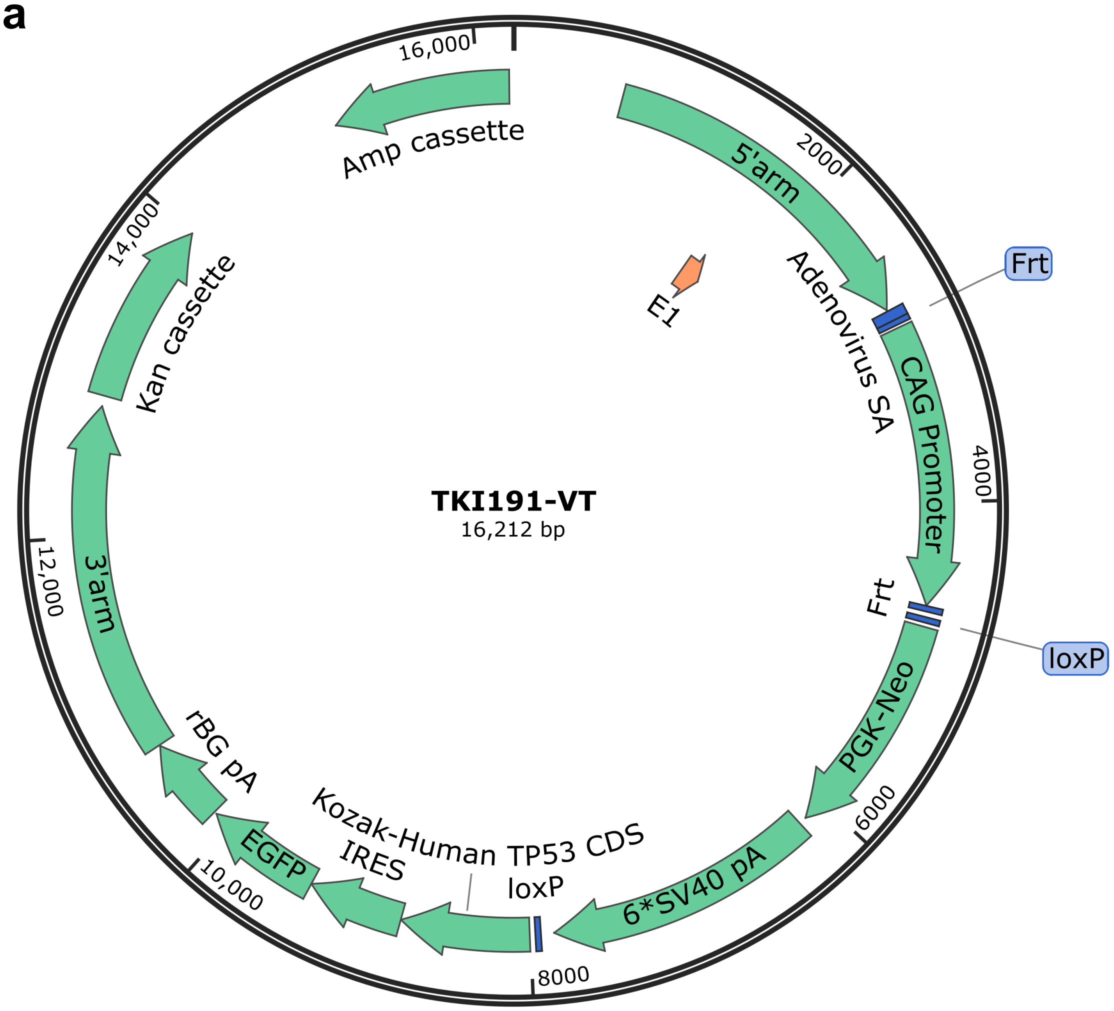
TKI191-VT
16,212 bp
16,000
2000
4000
6000
8000
10,000
12,000
14,000
Amp cassette
5'arm
Frt
E1
Adenovirus SA
CAG Promoter
Frt
loxP
PGK-Neo
6*SV40 pA
loxP
Kozak-Human TP53 CDS
IRES
EGFP
rBG pA
3'arm
Kan cassette

**b**

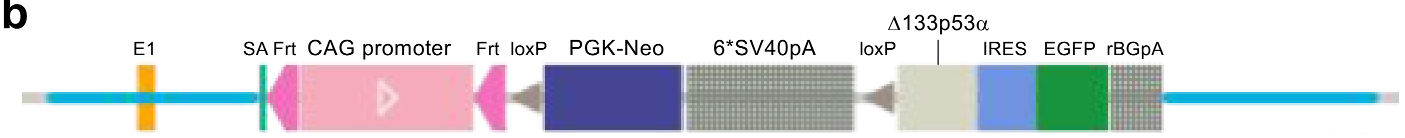
E1
SA
Frt
CAG promoter
Frt
loxP
PGK-Neo
6*SV40pA
loxP
Δ133p53α
IRES
EGFP
rBGpA

**Extended Data Fig. 1 | Targeting vector for generating Δ133p53α-transgenic mice. a,** Structure of the targeting vector for CAG-LSL-Δ133p53α allele (TKI191-VT). The elements are as follows (clockwise from the top): 5' arm homologous to the *ROSA26* locus; exon 1 (E1) of *ROSA26*; adenovirus splice acceptor (SA); Frt site (not used in this study); CAG promoter; Frt site (not used in this study); loxP site; PGK promoter-driven neomycin resistance gene (neo); six tandem repeats of SV40 poly(A) signals (6*SV40 pA, a transcription stop cassette); loxP site; Kozak-modified human *TP53* coding sequence (CDS) corresponding to Δ133p53α (NM_001126115.1); IRES; EGFP; rabbit β-globin poly(A) signal (rBG pA); 3' arm homologous to the *ROSA26* locus; Kan-resistance cassette; and Amp-resistance cassette. The full sequence of the vector is available upon request. **b,** Schematic representation of the *ROSA26* locus with the knocked-in targeting vector.

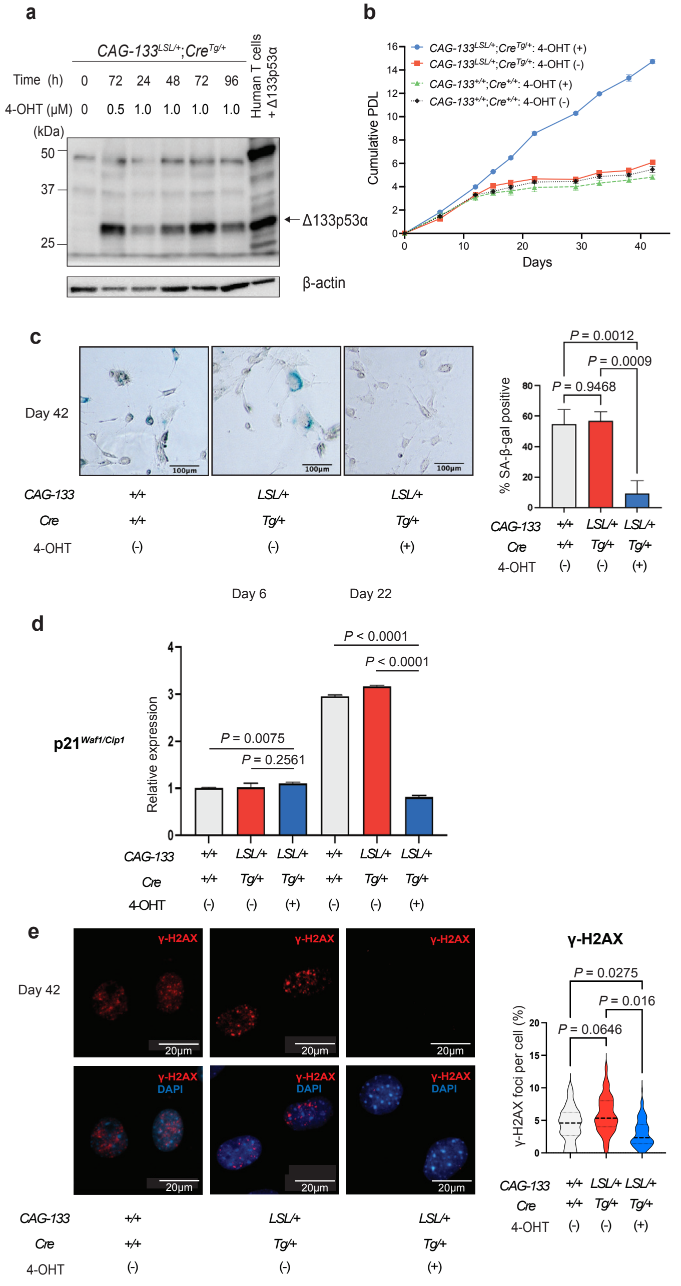
a
CAG-133LSL/+;CreTg/+
Human T cells + Δ133p53α
Time (h) 0 72 24 48 72 96
4-OHT (μM) 0 0.5 1.0 1.0 1.0 1.0
(kDa)
50
37
25
Δ133p53α
β-actin
b
CAG-133LSL/+;CreTg/+: 4-OHT (+)
CAG-133LSL/+;CreTg/+: 4-OHT (-)
CAG-133+/+;Cre+/+: 4-OHT (+)
CAG-133+/+;Cre+/+: 4-OHT (-)
Cumulative PDL
Days
c
Day 42
100μm
CAG-133 +/+ LSL/+ LSL/+
Cre +/+ Tg/+ Tg/+
4-OHT (-) (-) (+)
P = 0.0012
P = 0.0009
P = 0.9468
% SA-β-gal positive
Day 6
Day 22
d
p21Waf1/Cip1
Relative expression
P = 0.0075
P = 0.2561
P < 0.0001
P < 0.0001
CAG-133 +/+ LSL/+ LSL/+ +/+ LSL/+ LSL/+
Cre +/+ Tg/+ Tg/+ +/+ Tg/+ Tg/+
4-OHT (-) (-) (+) (-) (-) (+)
e
Day 42
γ-H2AX
DAPI
20μm
P = 0.0275
P = 0.016
P = 0.0646
γ-H2AX foci per cell (%)
CAG-133 +/+ LSL/+ LSL/+
Cre +/+ Tg/+ Tg/+
4-OHT (-) (-) (+)

**Extended Data Fig. 2 | Human Δ133p53α functions in mouse cells. a,** Western blot analysis showing induced expression of Δ133p53α in mouse embryonic fibroblasts (MEFs). MEFs with CAG-LSL-Δ133p53α and UBC-Cre-ERT2 alleles both in heterozygous states ($CAG\text{-}133^{LSL/+};Cre^{Tg/+}$) were treated with 0.5 or 1.0 μM 4-hydroxytamoxifen (4-OHT) for the indicated time periods. Human T cells with lentiviral expression of Δ133p53α[1] were used as a positive control. Δ133p53α, indicated by an arrow, was detected by a sheep polyclonal antibody SAPU. β-actin was a loading control. **b,** Cell proliferation of MEFs with and without Δ133p53α expression. $CAG\text{-}133^{LSL/+};Cre^{Tg/+}$ MEFs and wild-type MEFs ($CAG\text{-}133^{+/+};Cre^{+/+}$), untreated or treated with 1.0 μM 4-OHT for 72 h, were monitored for cell numbers, and their cumulative population doubling levels (PDL) were plotted to days in culture. **c,** Senescence-associated (SA)-β-galactosidase (gal) staining. Cells corresponding to those on day 42 in **b** were stained in biological triplicate. Representative images are shown on the left (scale bars, 100 μm). Quantitative summary of SA-β-gal-positive cells is on the right. Data are mean ± s.d. (n = 3), with at least 100 cells observed per biological replicate. *P* values were calculated using one-way ANOVA with Tukey's multiple comparison test. **d,** qRT-PCR analysis of $p21^{Waf1/Cip1}$ mRNA expression. Cells treated with 4-OHT corresponding to those on day 6 and day 22 in **b** were analyzed in biological triplicate. Data were normalized to GAPDH and expressed relative to wild-type MEFs on day 6, and are presented as mean ± s.d. (n = 3). *P* values were calculated using Welch's *t*-test. **e,** Immunofluorescence staining of γ-H2AX foci. Cells corresponding to those on day 42 in **b** were stained. Representative images are shown on the left (red, γ-H2AX; blue, DAPI; scale bars, 10 μm). On the right, all data from biological triplicate (n ≥ 100 cells per replicate) are presented as violin plots showing the number of γ-H2AX foci per cell, with a dashed bold line indicating the median and two thin lines indicating the first quartile and third quartile. *P* values were determined using nested one-way ANOVA.

1 Roselle, C. *et al.* Enhancing chimeric antigen receptor T cell therapy by modulating the p53 signaling network with Δ133p53α. *Proc. Natl. Acad. Sci. U. S. A.* **121**, e2317735121 (2024). https://doi.org:10.1073/pnas.2317735121

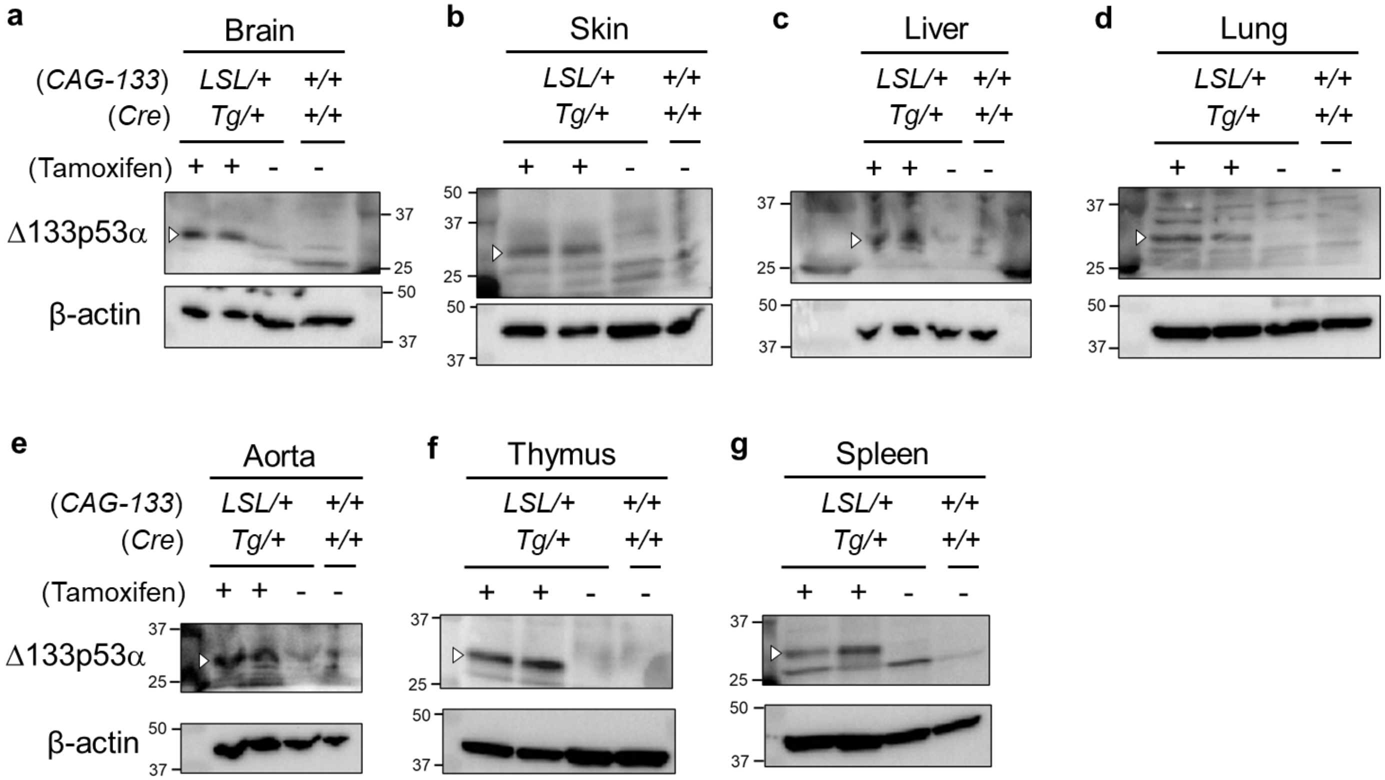
a
Brain
(CAG-133) LSL/+ +/+
(Cre) Tg/+ +/+
(Tamoxifen) + + - -
Δ133p53α
β-actin
37
25
50
37
b
Skin
LSL/+ +/+
Tg/+ +/+
+ + - -
50
37
25
50
37
c
Liver
LSL/+ +/+
Tg/+ +/+
+ + - -
37
25
50
37
d
Lung
LSL/+ +/+
Tg/+ +/+
+ + - -
37
25
50
37
e
Aorta
(CAG-133) LSL/+ +/+
(Cre) Tg/+ +/+
(Tamoxifen) + + - -
Δ133p53α
β-actin
37
25
50
37
f
Thymus
LSL/+ +/+
Tg/+ +/+
+ + - -
37
25
50
37
g
Spleen
LSL/+ +/+
Tg/+ +/+
+ + - -
37
25
50
37

**Extended Data Fig. 3 | Tamoxifen-induced expression of Δ133p53α in mouse tissues.**

Western blot detection of tamoxifen-induced transgenic expression of Δ133p53α (indicated by white arrowheads) in brain (**a**), skin (**b**), liver (**c**), lung (**d**), aorta (**e**), thymus (**f**) and spleen (**g**). The following 15-week-old mice were examined: two *CAG-133*$^{LSL/+}$;*Cre*$^{Tg/+}$ mice that had tamoxifen injection at 8-9 weeks of age; a *CAG-133*$^{LSL/+}$;*Cre*$^{Tg/+}$ mouse with no tamoxifen injection; and a *CAG-133*$^{+/+}$;*Cre*$^{+/+}$ mouse (wild-type control). Δ133p53α was detected by a sheep polyclonal antibody SAPU[2]. β-actin was a loading control. Size markers (kDa) are indicated.

2 Marcel, V., Khoury, M. P., Fernandes, K., Diot, A., Lane, D. P. & Bourdon, J. C. Detecting p53 isoforms at protein level. *Methods Mol. Biol.* **962**, 15-29 (2013). https://doi.org:10.1007/978-1-62703-236-0_2

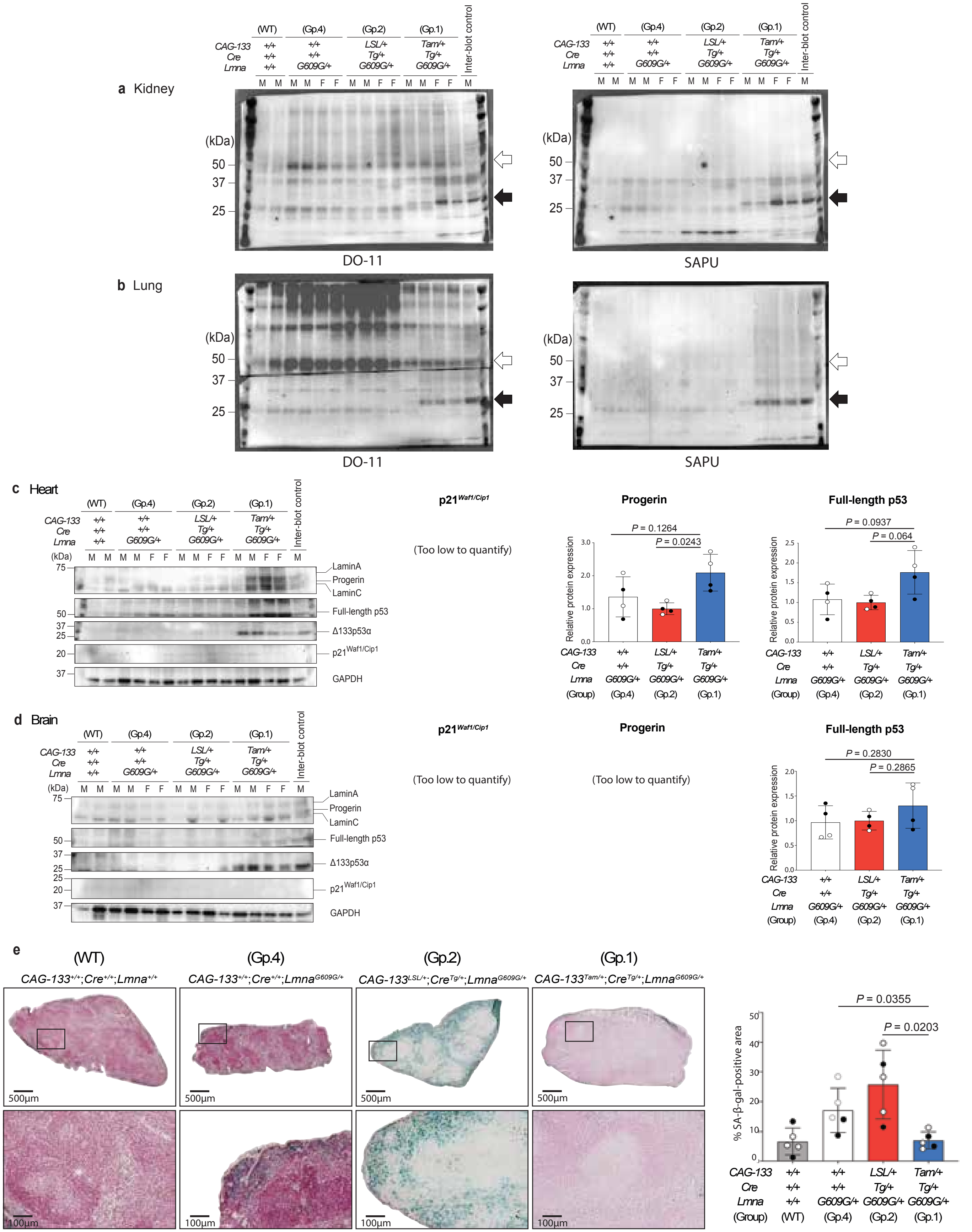

a Kidney
(WT)
(Gp.4)
(Gp.2)
(Gp.1)
Inter-blot control
CAG-133
Cre
Lmna
+/+
LSL/+
Tam/+
Tg/+
G609G/+
M
F
(kDa)
50
37
25
DO-11
SAPU
b Lung
DO-11
SAPU
c Heart
LaminA
Progerin
LaminC
Full-length p53
Δ133p53α
p21Waf1/Cip1
GAPDH
75
20
p21Waf1/Cip1
(Too low to quantify)
Progerin
P = 0.1264
P = 0.0243
Relative protein expression
Full-length p53
P = 0.0937
P = 0.064
(Group)
(Gp.4)
(Gp.2)
(Gp.1)
d Brain
Progerin
(Too low to quantify)
Full-length p53
P = 0.2830
P = 0.2865
e
(WT)
(Gp.4)
(Gp.2)
(Gp.1)
CAG-133+/+;Cre+/+;Lmna+/+
CAG-133+/+;Cre+/+;LmnaG609G/+
CAG-133LSL/+;CreTg/+;LmnaG609G/+
CAG-133Tam/+;CreTg/+;LmnaG609G/+
500µm
100µm
P = 0.0355
P = 0.0203
% SA-β-gal-positive area

**Extended Data Fig. 4 | Western blot and SA-β-gal assays in *Lmna*$^{G609G/+}$ progeria mice with and without Δ133p53α expression. a,b,** Western blot analysis using two anti-p53 antibodies, DO-11 (left) and SAPU (right), in the kidney (**a**) and lung (**b**). Single Western blot membranes, which contained protein samples from the same set of mice as in Fig. 1, were probed with DO-11 and SAPU, and the results are presented here side by side. The positions of full-length p53 (open arrows) and Δ133p53α (closed arrows) are indicated. Based on these results, DO-11 and SAPU were primarily used to detect full-length p53 and Δ133p53α, respectively, in this study. The cropped bands of full-length p53 and Δ133p53α shown in Fig. 1c (kidney) and Fig. 1e (lung) correspond to those detected here by DO-11 and SAPU, respectively, in **a** (kidney) and **b** (lung). **c,d,** Western blot analyses of lamin A/C and progerin, full-length p53, Δ133p53α, p21$^{Waf1/Cip1}$, and GAPDH (normalization control) in the heart (**c**) and brain (**d**). Protein samples were prepared from the same set of mice as in Fig. 1. The same inter-blot control as in Fig. 1 was included in both blots. Quantitative analysis and data presentation were performed as in Fig. 1 (mean ± s.d., n = 4, Welch's *t*-test). The expression levels of p21$^{Waf1/Cip1}$ in both the heart and brain (**c,d**) and of progerin in the brain (**d**) were too low to be quantified. **e,** SA-β-gal staining of the spleen. Spleen section slides from five 10-month-old mice per group, including Group 1 (*CAG-133*$^{Tam/+}$;*Cre*$^{Tg/+}$;*Lmna*$^{G609G/+}$), Group 2 (*CAG-133*$^{LSL/+}$;*Cre*$^{Tg/+}$;*Lmna*$^{G609G/+}$), Group 4 (*CAG-133*$^{+/+}$;*Cre*$^{+/+}$;*Lmna*$^{G609G/+}$), and wild-type control (*CAG-133*$^{+/+}$;*Cre*$^{+/+}$;*Lmna*$^{+/+}$), were stained and quantified as described in the Methods section. Representative images of the four groups are shown (scale bars; 500 μm in the whole sections above, 100 μm in the enlarged areas below). Quantitative data (% SA-β-gal-positive area) are presented as mean ± s.d. (n = 5; open circles, females; closed circles, males). *P* values were determined using Welch's *t*-test.

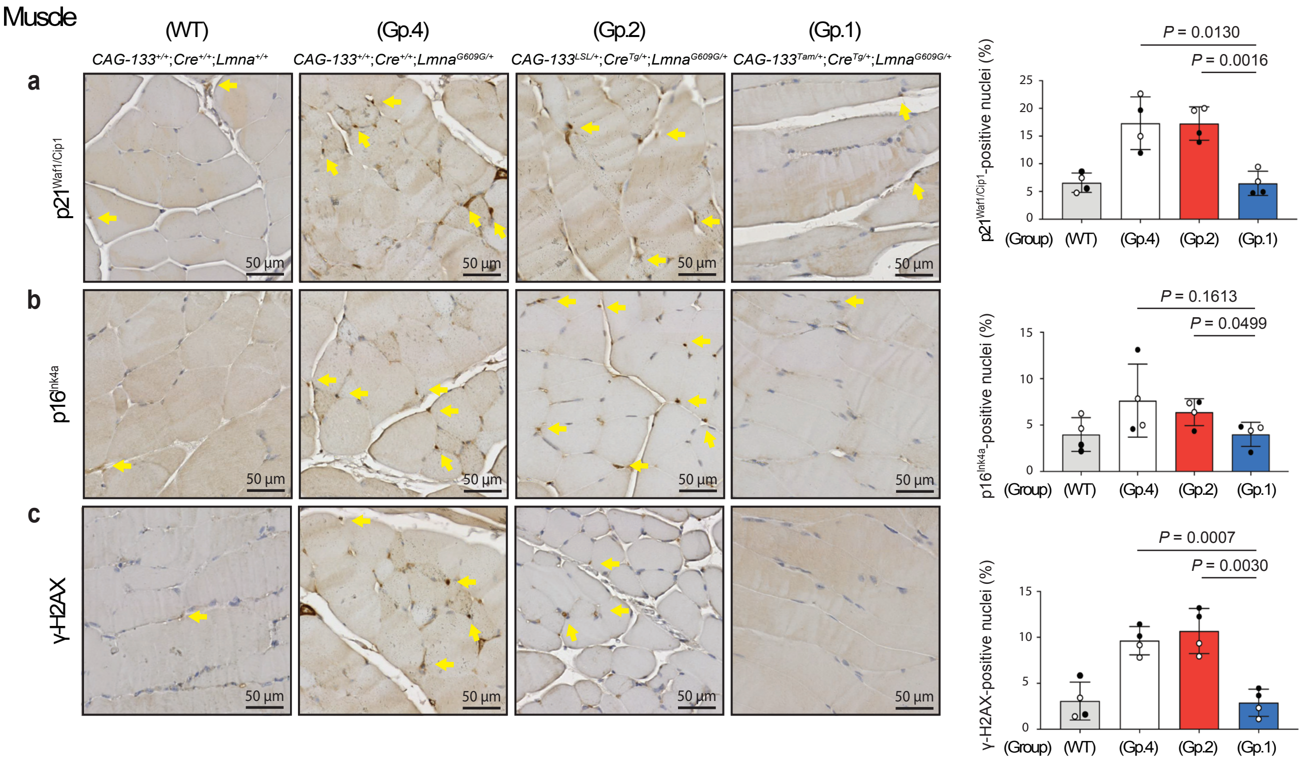

Muscle
(WT)
(Gp.4)
(Gp.2)
(Gp.1)
CAG-133+/+;Cre+/+;Lmna+/+
CAG-133+/+;Cre+/+;LmnaG609G/+
CAG-133LSL/+;CreTg/+;LmnaG609G/+
CAG-133Tam/+;CreTg/+;LmnaG609G/+
a
b
c
p21Waf1/Cip1
p16Ink4a
γ-H2AX
50 µm
P = 0.0130
P = 0.0016
p21Waf1/Cip1-positive nuclei (%)
P = 0.1613
P = 0.0499
p16Ink4a-positive nuclei (%)
P = 0.0007
P = 0.0030
γ-H2AX-positive nuclei (%)
(Group)

**Extended Data Fig. 5 | Cellular senescence and DNA damage reduced by Δ133p53α in the skeletal muscle. a-c,** IHC staining of the skeletal muscle from the same set of Group-1, Group-2, Group-4, and wild-type mice as in Fig. 2. Representative images of $p21^{Waf1/Cip1}$ (**a**), $p16^{Ink4a}$ (**b**), and γ-H2AX (**c**) are shown (scale bars, 50 μm). Yellow arrows indicate examples of positively stained nuclei. Quantitative summaries are presented on the right (mean ± s.d. from n = 4; open circles, females; closed circles, males). *P* values were determined by Welch's *t*-test.

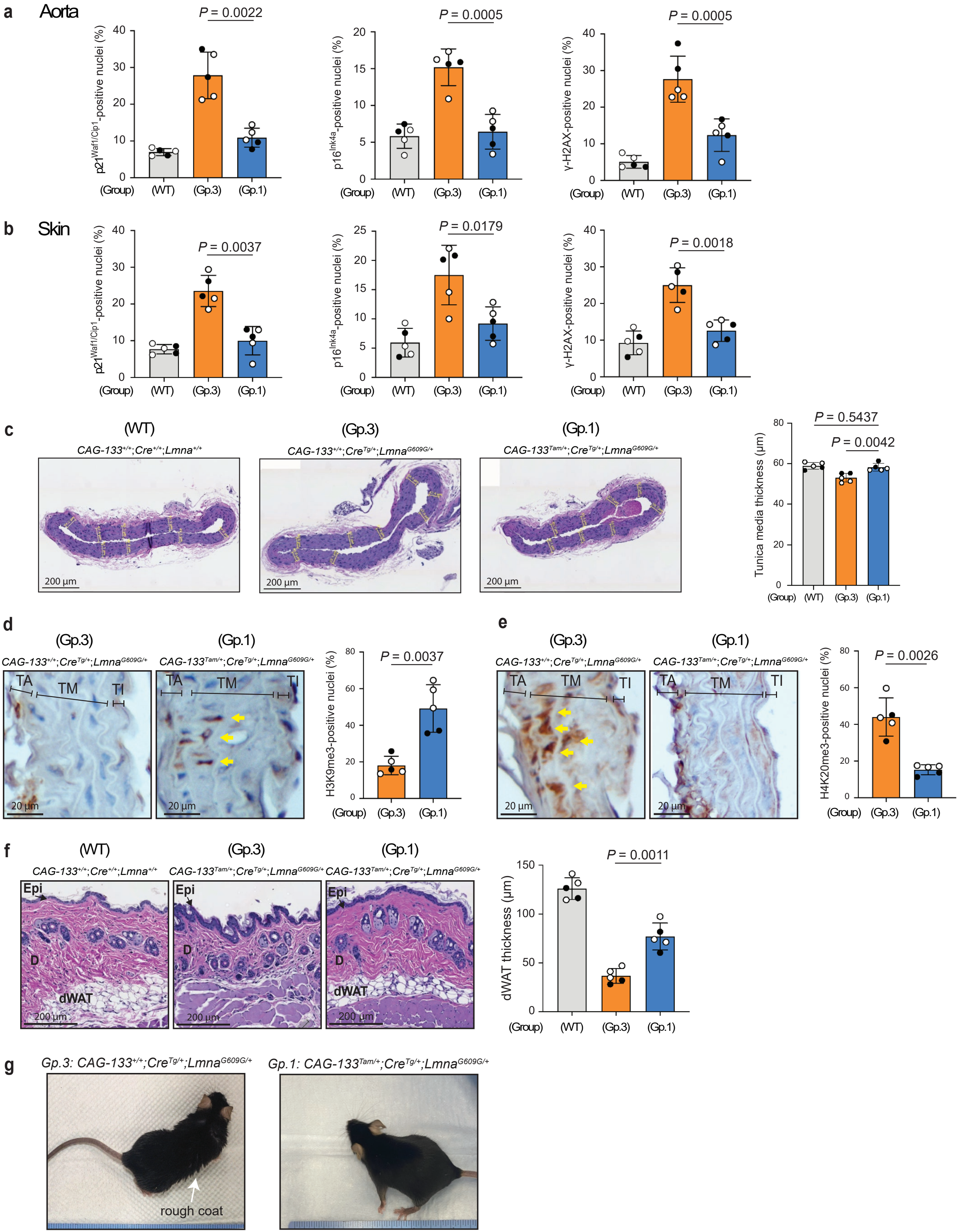

a Aorta
P = 0.0022
p21Waf1/Cip1-positive nuclei (%)
P = 0.0005
p16Ink4a-positive nuclei (%)
P = 0.0005
γ-H2AX-positive nuclei (%)
(Group) (WT) (Gp.3) (Gp.1)
b Skin
P = 0.0037
P = 0.0179
P = 0.0018
c
(WT) CAG-133+/+;Cre+/+;Lmna+/+
(Gp.3) CAG-133+/+;CreTg/+;LmnaG609G/+
(Gp.1) CAG-133Tam/+;CreTg/+;LmnaG609G/+
200 μm
P = 0.5437
P = 0.0042
Tunica media thickness (μm)
d
TA TM TI
20 μm
P = 0.0037
H3K9me3-positive nuclei (%)
e
P = 0.0026
H4K20me3-positive nuclei (%)
f
Epi D dWAT
P = 0.0011
dWAT thickness (μm)
g
Gp.3: CAG-133+/+;CreTg/+;LmnaG609G/+
Gp.1: CAG-133Tam/+;CreTg/+;LmnaG609G/+
rough coat

**Extended Data Fig. 6 | Tissue integrity in the aorta and skin rescued by Δ133p53α.** Tissue sections of the aorta and skin were from the same set of 9-10-month-old mice as in Fig. 3. **a,b,** Assessment of senescence and DNA damage markers. IHC staining of $p21^{Waf1/Cip1}$, $p16^{Ink4a}$, and γ-H2AX was performed and quantitatively analyzed as in Fig. 2. Quantitative summaries for the aorta (**a**) and skin (**b**) are presented as mean ± s.d. (n = 5; open circles, females; closed circles, males). *P* values were determined by Welch's *t*-test. **c,** Measurement of the thickness of the aortic tunica media. As shown in the representative images, thickness for each mouse was calculated as the average of measurements taken at eight randomly selected, approximately evenly spaced sites (indicated by yellow lines) Scale bars, 200 µm. Data are presented as mean ± s.d. (n = 5; open circles, females; closed circles, males). *P* values were determined using Welch's *t*-test. **d,e.** IHC staining of H3K9me3 (**d**) and H4K20me3 (**e**) in the aorta of Group-1 and Group-3 mice. Representative images (scale bars, 20 µm) show yellow arrows indicating examples of positively stained nuclei in the tunica media (TM). Quantitative data (% of positive nuclei) are presented as mean ± s.d. (n = 5; open circles, females; closed circles, males). *P* values were calculated using Welch's *t*-test. Non-specific staining is present in the tunica adventitia (TA) and tunica intima (TI). **f,** Measurement of the thickness of dermal white adipose tissue (dWAT) in the skin. The thickness (µm) for each mouse was calculated as the average of measurements taken at five randomly selected, approximately evenly spaced sites. Scale bars, 200 µm. Data are presented as mean ± s.d. (n = 5). *P* values were calculated using Welch's *t*-test. **g,** Photographs of Group-1 and Group-3 mice. Arrow indicates rough coat.

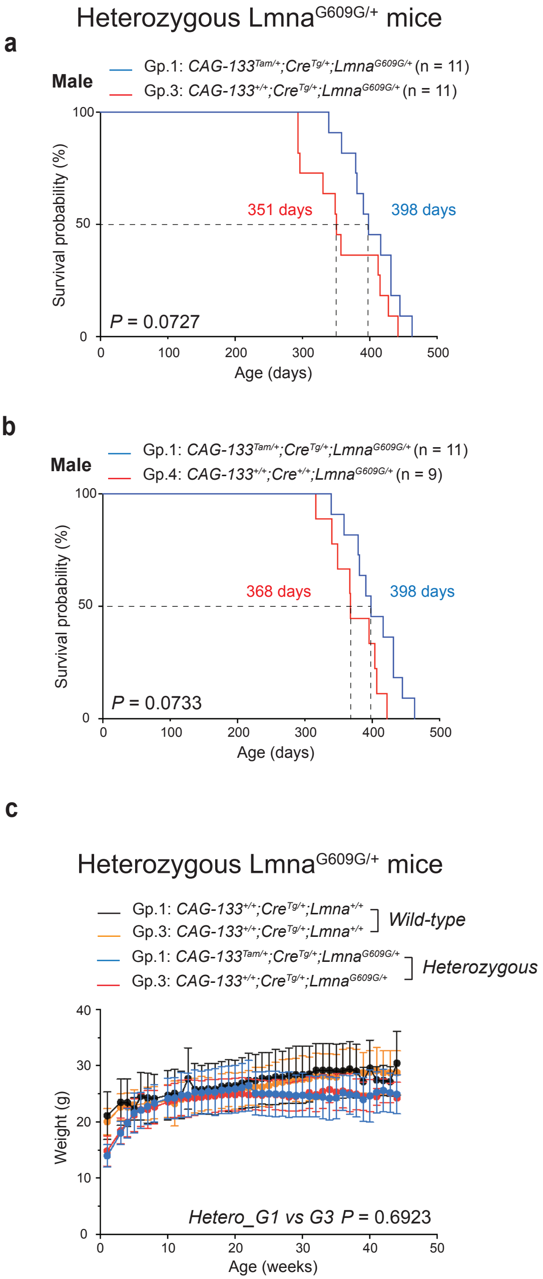
Heterozygous Lmna$^{G609G/+}$ mice
a
Male
Gp.1: CAG-133$^{Tam/+}$;Cre$^{Tg/+}$;Lmna$^{G609G/+}$ (n = 11)
Gp.3: CAG-133$^{+/+}$;Cre$^{Tg/+}$;Lmna$^{G609G/+}$ (n = 11)
Survival probability (%)
351 days
398 days
P = 0.0727
Age (days)
Female
Gp.1: CAG-133$^{Tam/+}$;Cre$^{Tg/+}$;Lmna$^{G609G/+}$ (n = 4)
Gp.3: CAG-133$^{+/+}$;Cre$^{Tg/+}$;Lmna$^{G609G/+}$ (n = 6)
330 days
385 days
P = 0.4485
b
Male
Gp.1: CAG-133$^{Tam/+}$;Cre$^{Tg/+}$;Lmna$^{G609G/+}$ (n = 11)
Gp.4: CAG-133$^{+/+}$;Cre$^{+/+}$;Lmna$^{G609G/+}$ (n = 9)
368 days
398 days
P = 0.0733
Female
Gp.1: CAG-133$^{Tam/+}$;Cre$^{Tg/+}$;Lmna$^{G609G/+}$ (n = 4)
Gp.4: CAG-133$^{+/+}$;Cre$^{+/+}$;Lmna$^{G609G/+}$ (n = 7)
344 days
385 days
P = 0.3695
c
Heterozygous Lmna$^{G609G/+}$ mice
Gp.1: CAG-133$^{+/+}$;Cre$^{Tg/+}$;Lmna$^{+/+}$
Gp.3: CAG-133$^{+/+}$;Cre$^{Tg/+}$;Lmna$^{+/+}$
Wild-type
Gp.1: CAG-133$^{Tam/+}$;Cre$^{Tg/+}$;Lmna$^{G609G/+}$
Gp.3: CAG-133$^{+/+}$;Cre$^{Tg/+}$;Lmna$^{G609G/+}$
Heterozygous
Weight (g)
Hetero_G1 vs G3 P = 0.6923
Age (weeks)

**Extended Data Fig. 7 | Monitoring *Lmna*$^{G609G/+}$ mice with and without Δ133p53α. a,b,** Kaplan-Meier survival curves comparing Group-1 mice with Group-3 mice (**a**) or Group-4 mice (**b**), when male and female mice were analyzed separately. Numbers of mice (n) are indicated. Median survival days and *P* values (log-rank test) are shown. **c,** Body weight charts of Group-1 and Group-3 *Lmna*$^{G609G/+}$ mice. Group-1 and Group-3 wild-type mice are shown for comparison. For each group, data from all surviving mice (mean ± s.d.) were plotted up to 44 weeks of age, at which point Group-3 *Lmna*$^{G609G/+}$ mice still retained three mice. No significant difference between Group-1 and Group-3 *Lmna*$^{G609G/+}$ mice ($P = 0.6923$, mixed-effects model).

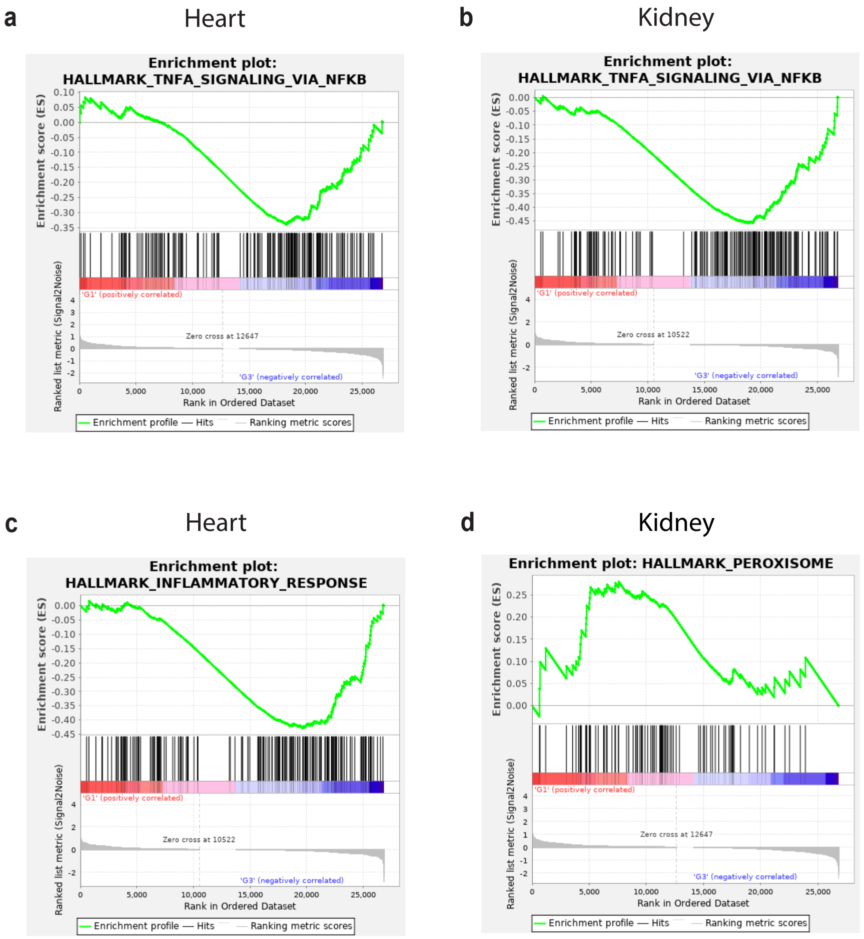
a
Heart
Enrichment plot: HALLMARK_TNFA_SIGNALING_VIA_NFKB
Enrichment score (ES)
Ranked list metric (Signal2Noise)
'G1' (positively correlated)
Zero cross at 12647
'G3' (negatively correlated)
Rank in Ordered Dataset
Enrichment profile — Hits — Ranking metric scores
b
Kidney
Enrichment plot: HALLMARK_TNFA_SIGNALING_VIA_NFKB
Enrichment score (ES)
Ranked list metric (Signal2Noise)
'G1' (positively correlated)
Zero cross at 10522
'G3' (negatively correlated)
Rank in Ordered Dataset
Enrichment profile — Hits — Ranking metric scores
c
Heart
Enrichment plot: HALLMARK_INFLAMMATORY_RESPONSE
Enrichment score (ES)
Ranked list metric (Signal2Noise)
'G1' (positively correlated)
Zero cross at 10522
'G3' (negatively correlated)
Rank in Ordered Dataset
Enrichment profile — Hits — Ranking metric scores
d
Kidney
Enrichment plot: HALLMARK_PEROXISOME
Enrichment score (ES)
Ranked list metric (Signal2Noise)
'G1' (positively correlated)
Zero cross at 12647
'G3' (negatively correlated)
Rank in Ordered Dataset
Enrichment profile — Hits — Ranking metric scores

**Extended Data Fig. 8 | Δ133p53α-induced changes in gene expression profiles revealed by RNA-seq.** Enrichment plots (Group 1 versus Group 3) are presented as in Fig. 5c-g. **a,b,** Downregulation of the TNFα signaling via NFkB in the heart (**a**) and kidney (**b**). **c,** Downregulation of the inflammatory response in the heart. **d,** Upregulation of the peroxisome pathway in the kidney. All leading edge genes in each pathway are listed in Extended Data Table 2.

**a** Homozygous Lmna$^{G609G/G609G}$ mice

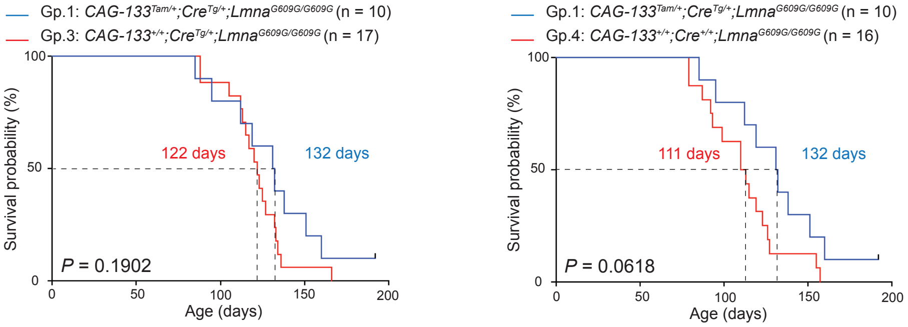


**b** Homozygous Lmna$^{G609G/G609G}$ mice

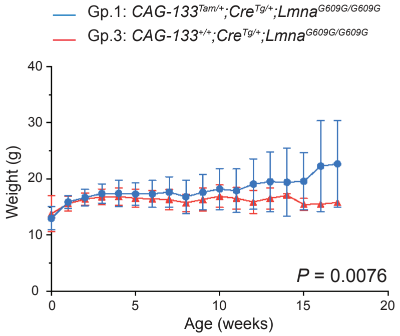

**Extended Data Fig. 9 | Lifespan and body weight of homozygous *Lmna*$^{G609G/G609G}$ mice with and without Δ133p53α. a,** Kaplan-Meier survival curves comparing homozygous Group-1 mice (*CAG-133*$^{Tam/+}$;*Cre*$^{Tg/+}$;*Lmna*$^{G609G/G609G}$) (n = 10) with Group-3 mice (*CAG-133*$^{+/+}$;*Cre*$^{Tg/+}$;*Lmna*$^{G609G/G609G}$) (n = 17, left) or Group-4 mice (*CAG-133*$^{+/+}$;*Cre*$^{+/+}$;*Lmna*$^{G609G/G609G}$) (n = 16, right). Both female and male mice were included. Median survival days and *P* values (log-rank test) are indicated. **b,** Body weight charts of Group-1 and Group-3 *Lmna*$^{G609G/G609G}$ mice. Data from all surviving mice (mean ± s.d.) were plotted up to 17 weeks of age. *P* = 0.0076 (mixed-effects model).
